# Smoothened Transduces Hedgehog Signals via Activity-Dependent Sequestration of PKA Catalytic Subunits

**DOI:** 10.1101/2020.07.01.183079

**Authors:** Corvin D. Arveseth, John T. Happ, Danielle S. Hedeen, Ju-Fen Zhu, Jacob L. Capener, Dana Klatt Shaw, Ishan Deshpande, Jiahao Liang, Jiewei Xu, Sara L. Stubben, Isaac B. Nelson, Madison F. Walker, Nevan J. Krogan, David J. Grunwald, Ruth Hüttenhain, Aashish Manglik, Benjamin R. Myers

**Author notes:** Co-first author (Corvin D. Arveseth, John T. Happ). Co-second author (Danielle S. Hedeen, Ju-Fen Zhu). Department of Developmental Biology, Washington University School of Medicine, St. Louis, MO 63110 USA. Department of Plant and Wildlife Sciences, College of Life Sciences, Brigham Young University, Provo, UT 84602 USA.

## Abstract

The Hedgehog (Hh) pathway is essential for organ development, homeostasis, and regeneration. Dysfunction of this cascade drives several cancers. To control expression of pathway target genes, the G protein-coupled receptor (GPCR) Smoothened (SMO) activates glioma-associated (GLI) transcription factors via an unknown mechanism. Here we show that, rather than conforming to traditional GPCR signaling paradigms, SMO activates GLI by binding and sequestering protein kinase A (PKA) catalytic subunits at the membrane. This sequestration, triggered by GPCR kinase 2 (GRK2)-mediated phosphorylation of SMO intracellular domains, prevents PKA from phosphorylating soluble substrates, releasing GLI from PKA-mediated inhibition. Our work provides a mechanism directly linking Hh signal transduction at the membrane to GLI transcription in the nucleus. This process is more fundamentally similar between species than prevailing hypotheses suggest. The mechanism described here may apply broadly to other GPCR- and PKA-containing cascades in diverse areas of biology.

## INTRODUCTION

The Hedgehog (Hh) pathway controls the development of nearly every vertebrate organ (Briscoe and Thérond, 2013; Ingham and McMahon, 2001; Ingham et al., 2011; Kong et al., 2019). It also plays critical roles in stem cell biology and injury-induced tissue regeneration (Petrova and Joyner, 2014; Roberts et al., 2017). Insufficient pathway activation during embryogenesis gives rise to birth defects (Muenke and Beachy, 2000), whereas ectopic pathway activity drives several malignancies, including basal cell carcinoma of the skin and pediatric medulloblastoma (Pak and Segal, 2016; Wu et al., 2017).

Hh signal reception at the membrane is tightly coupled to transcriptional regulation of pathway target genes in the nucleus (Briscoe and Thérond, 2013; Kong et al., 2019; Kozielewicz et al., 2020; Qi and Li, 2020). In the pathway “off” state, Patched1 (PTCH1) inhibits the G protein-coupled receptor (GPCR) Smoothened (SMO). In the pathway “on” state, Hh proteins bind to and inactivate PTCH1, relieving SMO from inhibition (Briscoe and Thérond, 2013; Kong et al., 2019; Kozielewicz et al., 2020; Qi and Li, 2020). SMO activation ultimately results in the conversion of glioma-associated (GLI) transcription factors from repressor to activator forms (Briscoe and Thérond, 2013; Kong et al., 2019; Qi and Li, 2020). Active GLI regulates expression of Hh pathway target genes that drive cell differentiation or proliferation (Hui and Angers, 2011). The process by which vertebrate SMO activates GLI, however, is largely a mystery.

An appealing model suggests that SMO activates GLI by blocking protein kinase A (PKA), thereby releasing GLI from PKA-mediated inhibition (Alcedo et al., 1996; Ayers and Thérond, 2010; Heuvel and Ingham, 1996; Kong et al., 2019). In support of this model, inactivation of PKA catalytic subunits (PKA-C) induces the Hh pathway to near-maximal levels (Hammerschmidt et al., 1996; Huang et al., 2002; Jiang and Struhl, 1995; Li et al., 1995; Tuson et al., 2011). In addition, PKA phosphorylation of GLI hinders its transcriptional activity, while SMO activation results in loss of phosphorylation at these sites (Aza-Blanc et al., 1997; Humke et al., 2010; Méthot and Basler, 1999; Niewiadomski et al., 2013; Wang et al., 2000). Furthermore, SMO, PKA, and GLI may communicate directly with one another within a cell surface organelle known as the primary cilium, as all three proteins localize in or near this subcellular compartment (Barzi et al., 2009; Corbit et al., 2005; Gigante and Caspary, 2020; Haycraft et al., 2005; Kim et al., 2009; Rohatgi et al., 2007; Tuson et al., 2011). Nevertheless, the above model is controversial because G proteins, which canonically link GPCR activation to PKA inhibition, are not required for SMO to activate GLI (Low et al., 2008; Regard et al., 2013; Riobo et al., 2006). Thus, although PKA has been implicated in communication between SMO and GLI, the molecular mechanism by which SMO activates GLI remains poorly understood (Ayers and Thérond, 2010; Briscoe and Thérond, 2013; Kong et al., 2019; Qi and Li, 2020).

To dissect SMO-GLI communication, we used a heterologous cellular system to identify and reconstitute the Hh pathway step immediately downstream of SMO. We then characterized the underlying biochemical mechanism, and assessed the physiological relevance of our findings using established cellular and embryological assays of Hh signal transduction. Using this approach, we found that activated SMO blocks PKA substrate phosphorylation by directly binding and sequestering PKA-C subunits at the membrane. This prevents PKA phosphorylation of GLI, thereby triggering GLI activation. PKA-C binding to SMO is controlled by GRK family kinases that selectively phosphorylate the SMO active conformation on conserved residues in the intracellular domain. Our work reveals an unconventional route by which GPCRs can control PKA activity – one that may also be utilized by other signaling pathways that employ these proteins.

## RESULTS

### SMO inhibits PKA substrate phosphorylation in a G protein-independent manner

We hypothesized that SMO can inhibit PKA via a G protein-independent process. To test this hypothesis, we set up a model system to study SMO regulation of PKA. GLI-based readouts are problematic in this regard, as they are affected by manipulation of either SMO or PKA; this makes it difficult to determine whether SMO and PKA reside in the same linear pathway or constitute two separate influences that converge on GLI. To overcome this and other confounding factors (see “Supplemental Information”), we reconstituted SMO regulation of PKA in a HEK293 model system using a non-GLI readout of PKA activity. This approach also allows us to employ CRISPR, biochemical, and fluorescence-based tools that are uniquely robust in HEK293 cells.

To report PKA activity, we utilized CREB (cyclic AMP response element (CRE) binding protein) transcription factors **(Figure 1A)**. CREB is activated by PKA phosphorylation (Shaywitz and Greenberg, 1999), but is not known to be subject to the other major mechanisms that regulate GLI activity (Hui and Angers, 2011; Shaywitz and Greenberg, 1999). Our studies employed C-terminally-truncated versions of SMO (either SMO657 or SMO674, **Figure 1—figure supplement 1A,B**). These truncations contain the proximal segment of the cytoplasmic tail (pCT) that is essential for GLI activation but lack the nonessential distal segment (dCT) **(Figure 1—figure supplement 1A,C)** (Kim et al., 2015; Varjosalo et al., 2006). Removing the dCT improves SMO expression levels and detergent solubility (data not shown), thereby facilitating our subsequent biochemical analyses.

**Figure 1:**
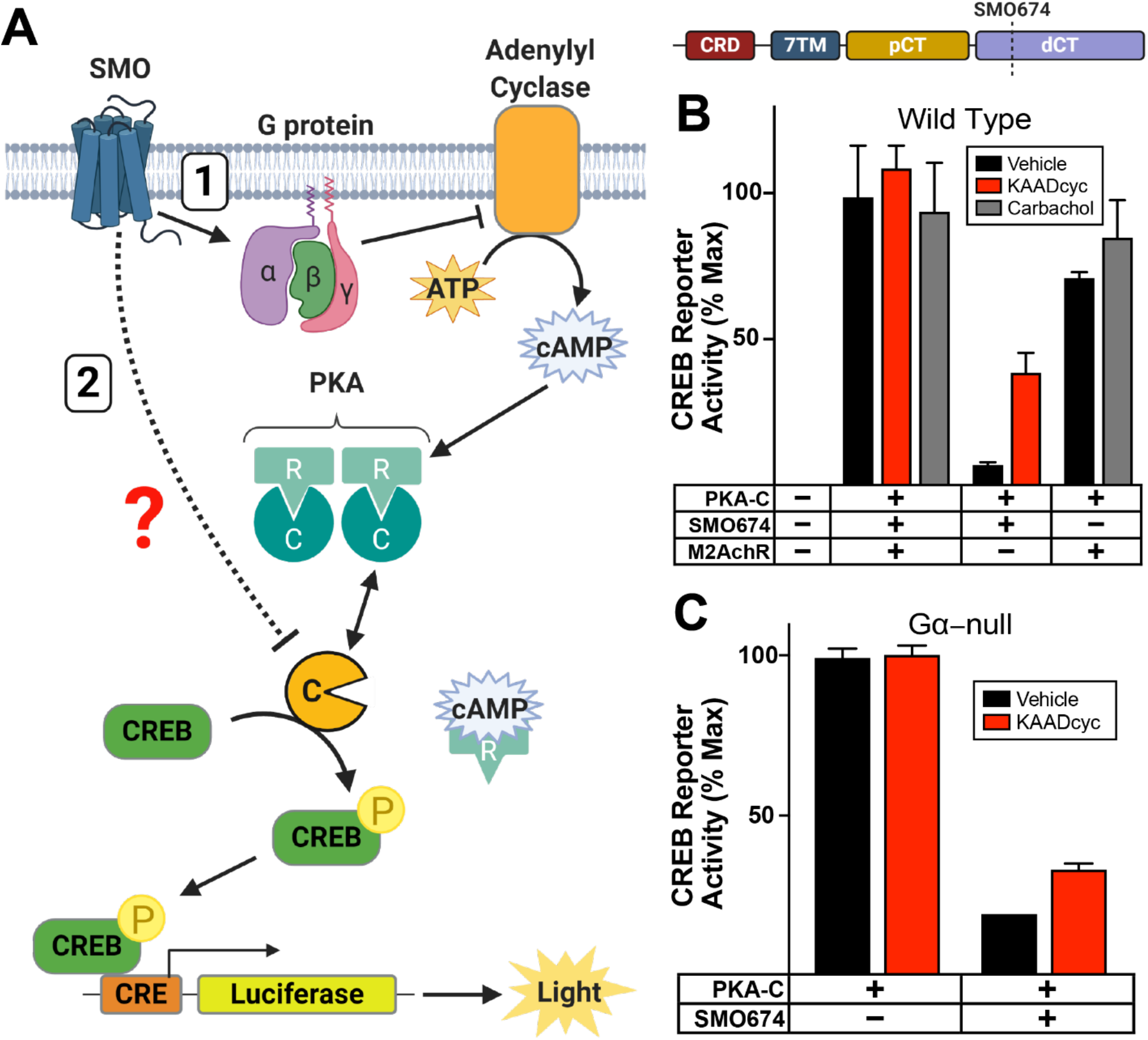
SMO inhibits PKA substrate phosphorylation in a G protein-independent manner. (**A**) Schematic of assay to detect phosphorylation of soluble PKA substrates. PKA-C phosphorylates CREB which binds CRE, inducing expression of luciferase. SMO can inhibit PKA-C by decreasing cAMP via inhibitory G proteins and Adenylyl Cyclase (AC) (route “1”). Alternatively, SMO may inhibit PKA-C via a G protein-independent mechanism (route “2”). (**B**) Wild-type HEK293 cells were transfected with CRE-luciferase reporter plasmid and GFP (as a negative control) or PKA-C, either alone, with SMO674 (see cartoon above), or with a canonical Gα_i/o_-coupled GPCR, M2AchR. Transfected cells were treated with the indicated drugs (vehicle control, M2AchR ligand carbachol (3 μM), or SMO antagonist KAADcyc (1 μM)). Following drug treatment, cells were lysed and luminescence measured. Note that transfected SMO is constitutively active in HEK293 cells because its inhibitor PTCH1 is present at minimal levels (Masdeu et al., 2006; Myers et al., 2017; Riobo et al., 2006; Shen et al., 2013), whereas M2AchR requires carbachol for activity. For the sake of clarity, the SMO constructs utilized in each experiment are indicated in the corresponding figure panel. (See Figure 1—figure supplement 1B for additional information.) (**C**) HEK293 Gα-null cells were transfected with PKA-C, either alone or with SMO674, and treated with vehicle or KAADcyc (1 μM). Data are normalized to 100%, which reflects reporter activation from PKA-C-transfected cells treated with vehicle (n = 3 biological replicates per condition, error bars = s.e.m.). See Supplemental Table 1 for statistical analysis.

In its active state, SMO, like many GPCRs, can block PKA-C by engaging inhibitory G proteins (Gα_i/o/z_) that inactivate adenylyl cyclase (AC), decrease cyclic AMP (cAMP), and promote PKA-C binding to regulatory (PKA-R) subunits to form an inactive holoenzyme **(Figure 1A, route “1”)**. We hypothesized, however, that SMO may directly inhibit free PKA-C subunits via a G protein-independent mechanism **(Figure 1A, route “2”)**. In this case, active SMO, but not other Gα_i/o/z_-coupled GPCRs, would block CREB reporter activation mediated by G protein-independent pathways.

To test this hypothesis, we expressed exogenous PKA-C, at levels likely to exceed those of endogenous PKA-R, to bypass G protein-dependent cascades. As expected, PKA-C expression strongly activated the CREB reporter **(Figure 1B)**, indicating an excess of PKA-C over PKA-R.

Cotransfection of SMO blocked PKA-C-mediated reporter activation **(Figure 1B)**, indicating that SMO can inhibit PKA-C in a G protein-independent manner. (Note that SMO is constitutively active in HEK293 cells, as its inhibitor PTCH1 is expressed at minimal levels (DeCamp et al., 2000; Masdeu et al., 2006; Myers et al., 2017; Riobo et al., 2006; Shen et al., 2013)). This blockade was partially reversed by the specific SMO antagonist KAAD-cyclopamine (KAADcyc), indicating that it depends on SMO activity. KAADcyc completely reversed effects of SMO in experiments where SMO blocked the reporter submaximally (data not shown). In contrast, activation of a canonical Gα_i/o_-coupled GPCR, the M2 acetylcholine receptor (M2AchR), with its ligand carbachol did not block the effects of PKA-C expression **(Figure 1B)**. This result cannot be explained by issues with receptor expression, trafficking, or ligand stimulation, because carbachol treatment of M2AchR-expressing cells readily blocked AC-evoked reporter activation **(Figure 1—figure supplement 1D)**. These experiments indicate that SMO can regulate PKA-C in a G protein-independent manner.

We verified this conclusion by showing that SMO also blocked PKA-C in HEK293 Gα-null cells harboring CRISPR-mediated deletions in all 13 human Gα genes (Hisano et al., 2019) **(Figure 1C)**. In contrast, M2AchR did not **(Figure 1—figure supplement 1D)**, consistent with its function as a canonical Gα_i/o_-coupled GPCR. Taken together, our findings indicate that SMO can inhibit PKA substrate phosphorylation even when G proteins are absent.

### SMO uses its essential pCT domain to recruit PKA-C to the membrane

GPCRs control PKA in part via G protein and cAMP-dependent regulation of its enzymatic activity (Taylor et al., 2012, 2013). However, another critical determinant of substrate phosphorylation is PKA’s localization within the cell. Interactions of PKA with specific receptors, signaling scaffolds, and anchoring proteins can bias its enzymatic activity towards certain subcellular locations and away from others (Scott and Pawson, 2009; Torres-Quesada et al., 2017). We hypothesized that SMO might control PKA subcellular localization, thereby restricting access of PKA to soluble substrates. Such a model could explain how SMO blocks CREB phosphorylation without requiring G proteins **(Figure 1)**.

We tested this hypothesis by examining the effect of SMO on the subcellular localization of PKA-C in HEK293 cells. In the presence of SMO, PKA-C localized to the membrane, colocalizing with SMO in a majority of cells (**Figure 2A,C**). In contrast, PKA-C did not display this membrane localization when SMO was absent (data not shown). SMO colocalized with PKA-C to a similar extent as it did with the nanobody (Nb) NbSmo2; this Nb binds efficiently and specifically to the activated conformation of SMO (**Figure 2A, Figure 2—figure supplement 1A,B, and Figure 2–figure supplement 2)** via an intracellular epitope within the seven-transmembrane (7TM) domain **(Figure 2—figure supplement 1C,D)**. As a control, the non-SMO-binding Nb, Nbβ2AR80 (Irannejad et al., 2013; Rasmussen et al., 2011), did not colocalize with SMO **(Figure 2A and Figure 2–figure supplement 2)**. Deletion of the SMO pCT abolished colocalization with PKA-C, but not with NbSmo2 **(Figure 2B,C and Figure 2–figure supplement 2)**. Thus, the SMO pCT, which is essential for activation of GLI **(Figure 1–figure supplement 1C)**, is also required to sequester PKA-C at the membrane.

**Figure 2:**
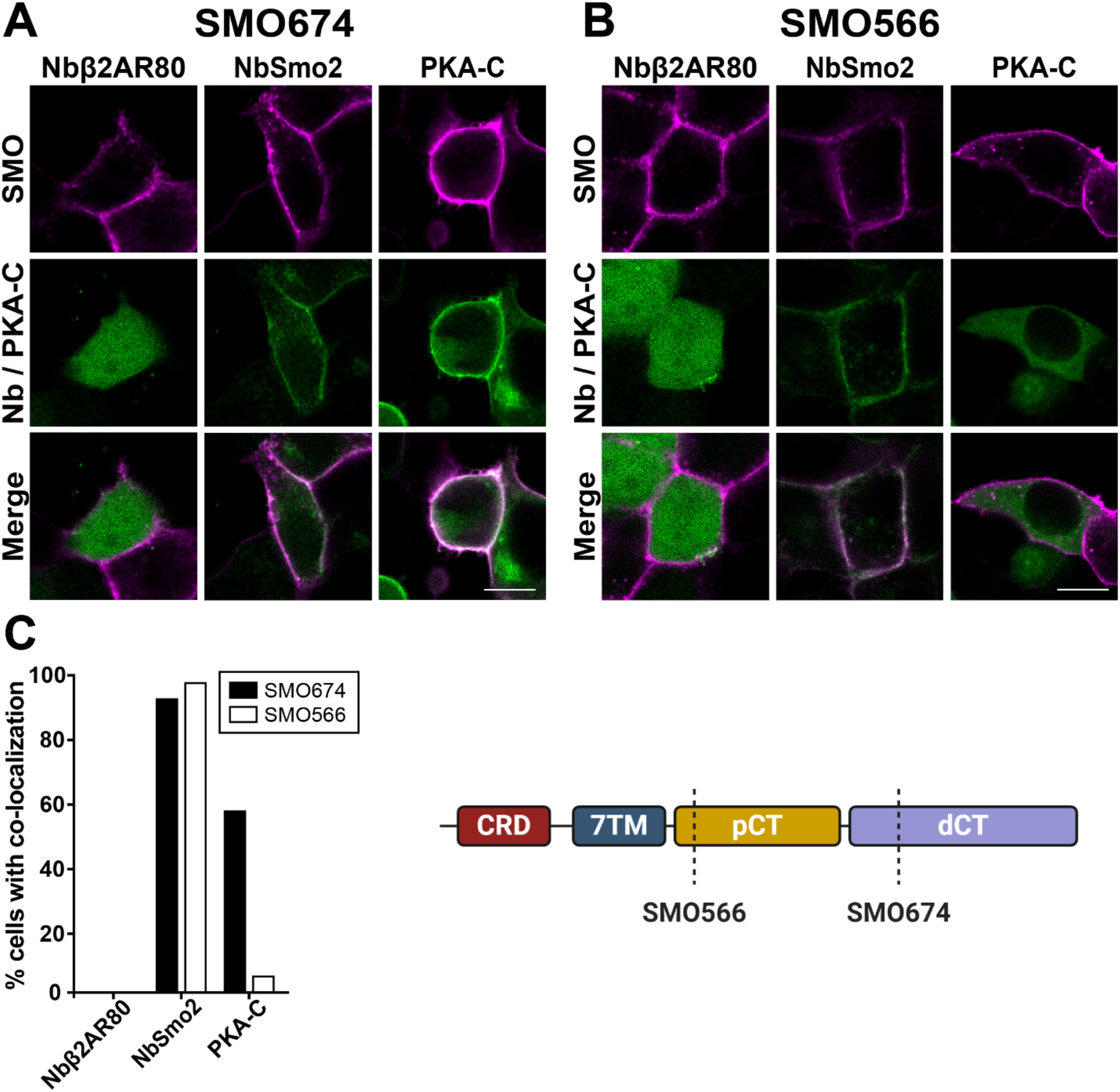
SMO uses its essential pCT domain to recruit PKA-C to the membrane. Cells expressing (**A**) FLAG-tagged SMO674 that contains the pCT or (**B**) FLAG-tagged SMO566 that lacks the pCT (see cartoon at lower right), were co-transfected with Nbβ2AR80-GFP, NbSmo2-YFP or PKA-C-YFP. Confocal microscopy images show SMO (magenta) and co-expressed proteins (green). (**C**) Percent of transfected cells that displayed colocalization in (A) and (B). Scale bar = 10 μm. (n = 29-121 cells per condition).

### The SMO pCT interacts with PKA-C

Vertebrate SMO may recruit PKA-C to the membrane **(Figure 2)** via a direct protein-protein interaction. Consistent with this hypothesis, *Drosophila* Smo associates with PKA-C subunits (Li et al., 2014; Ranieri et al., 2014). A comparable interaction, however, has not been reported for vertebrate SMO. To allow sensitive detection of protein-protein interactions in living cells without solubilization or wash steps that can disrupt labile interactions in conventional biochemical assays, we used bioluminescence resonance energy transfer (BRET) (Marullo and Bouvier, 2007). We fused SMO to a luciferase energy donor (nanoluc) and PKA-C or other candidate interactors to a YFP acceptor. Upon interaction (i.e, within ~10 nm of SMO), light produced by luciferase excites YFP **(Figure 3A, left)**. The YFP/luciferase emission ratio thus provides a normalized metric for protein interactions with SMO.

**Figure 3:**
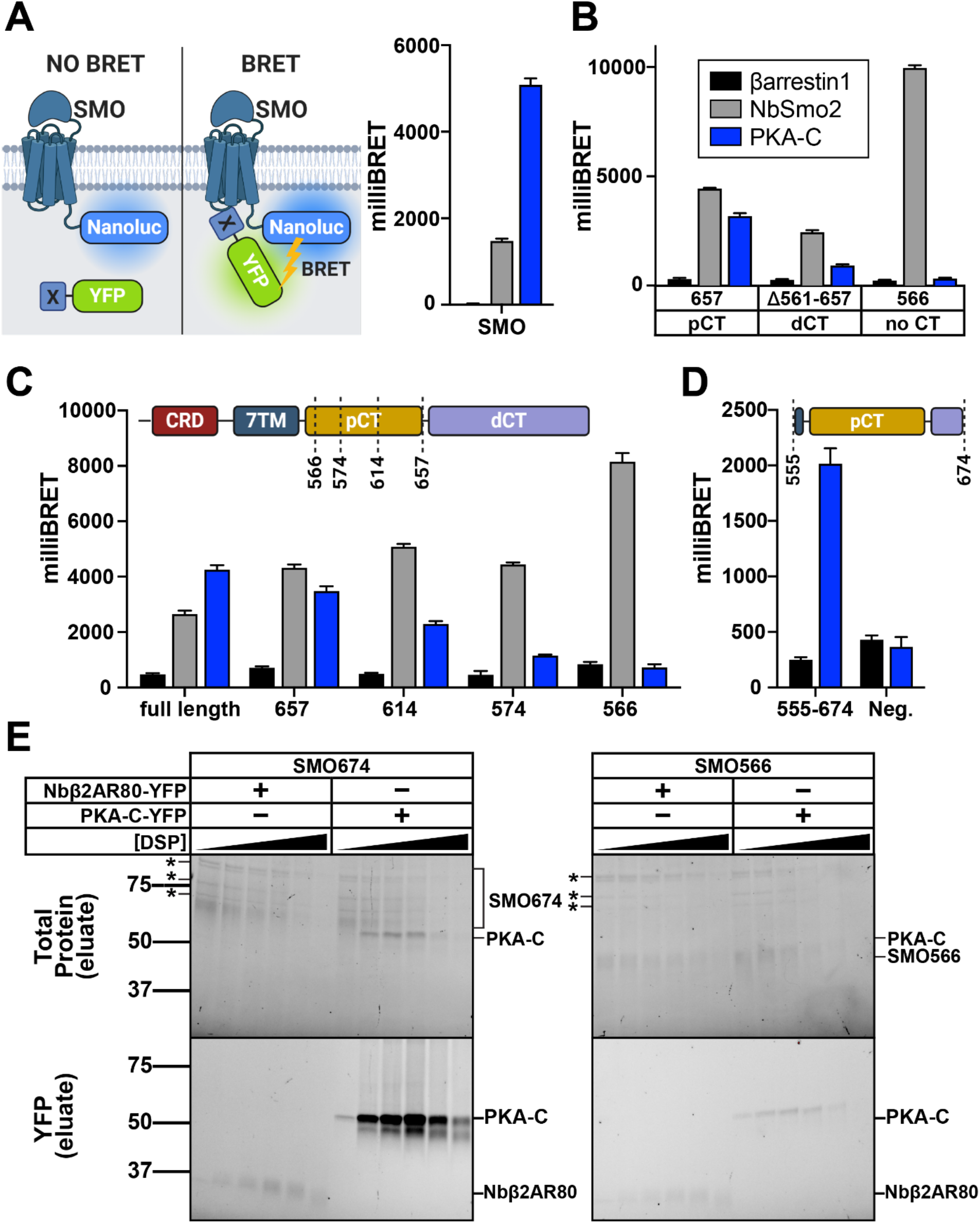
The SMO pCT interacts with PKA-C. (**A**) (Left) Schematic showing BRET between a nanoluc-tagged donor (SMO-nanoluc) and a YFP-tagged acceptor. For all BRET experiments in this figure, HEK293 cells were transfected with nanoluc-tagged SMO, along with YFP-tagged βarrestin-1 (black), NbSmo2 (gray) or PKA-C (blue). (Right) BRET experiment employing SMO with a full-length cytoplasmic tail. (**B**) Nanoluc-tagged SMO657 (which contains the pCT), SMOΔ561-657 (which contains the dCT), or SMO566 (which lacks the CT entirely) serve as donors. Note that NbSmo2 binding does not require the SMO CT (see Figure 2—figure supplement 1). In fact, NbSmo2 BRET increases upon SMO CT truncation, likely because the decreased distance between the NbSmo2 binding site and the nanoluc tag leads to more efficient BRET. (**C**) Nanoluc fusions of successive SMO CT truncations (SMO, SMO657, SMO614, SMO574, and SMO566) were utilized to determine the region of the pCT required for efficient BRET with PKA-C. Cartoon above the graph indicates the position of each CT truncation. (**D**) BRET between PKA-C and a Protein C- and nanoluc-tagged SMO pCT construct, without extracellular or 7TM domains (SMO555-674, see cartoon above). The same construct lacking SMO sequences (“Neg.”) serves as a negative control. (**E**) HEK293 cells were infected with viruses encoding FLAG-tagged SMO674 or SMO566, and YFP-tagged PKA-C or Nbβ2AR80, and treated with increasing concentrations of DSP crosslinker (0, 0.125, 0.25, 0.5, 1, or 2 mM). Following DSP quenching, cell lysis, and FLAG purification of SMO complexes, purified samples were separated on reducing SDS-PAGE. Total protein (top) and in-gel YFP fluorescence scans (bottom) for FLAG eluates are shown. * = copurifying contaminant proteins. Molecular masses are in kDa. Recovery of SMO / PKA-C complexes declines at DSP concentrations above 1 mM, likely because high DSP concentrations induce protein aggregation which decreases soluble protein yields in total cell lysates (see Figure 3–figure supplement 2). Data are reported as milliBRET ratios (YFP/Renilla x 1000), and background BRET values derived from cells expressing SMO-nanoluc alone were subtracted from all measurements. (n = 3-6 biological replicates per condition. Error bars = s.e.m.) See Supplemental Table 1 for statistical analysis.

SMO and PKA-C produced extremely strong BRET signals that often exceeded those of our NbSmo2 positive control **(Figure 3A, right)**. SMO BRET with PKA-C mainly required the pCT but not the dCT (compare SMO657 to SMOΔ561-657) **(Figure 3B)**. As expected, SMO BRET with NbSmo2 was efficient regardless of the CT **(Figure 3B)**. Truncations within the SMO pCT revealed that amino acids 574 to 657 are crucial for BRET with PKA-C **(Figure 3C and Figure 1–figure supplement 1B)**. SMO and PKA-C therefore interact in living cells in a pCT-dependent manner.

The BRET between SMO and PKA-C reflects a *bona fide* protein-protein interaction, based on several observations. First, titration of a fixed amount of SMO against a varying amount of PKA-C revealed a saturable BRET response **(Figure 3—figure supplement 1A)**, indicative of specific binding rather than nonspecific crowding-induced collisions. Second, a control membrane protein, PTCH1, fails to exhibit BRET with PKA-C **(Figure 3—figure supplement 1B)**. Several other proteins, including Nbβ2AR80, suppressor of Fused (SUFU) (Kong et al., 2019), and βarrestins (Shenoy and Lefkowitz, 2011), also showed minimal or no BRET with SMO **(Figure 3A and Figure 3—figure supplement 1C,D)**. Third, BRET is specific, since it requires defined sequences within SMO **(Figure 3B,C)**. Finally, BRET is unlikely an artifact of protein overexpression, as BRET signals do not correlate with expression levels of acceptor proteins **(Figure 3—figure supplement 1E)**.

To test whether the SMO pCT is sufficient to interact with PKA-C, we studied a soluble SMO construct containing this domain but lacking the extracellular and seven-transmembrane (7TM) regions (SMO555-674). This construct showed BRET with PKA-C **(Figure 3D)**, albeit at lower levels than SMO containing the 7TM domain (compare to **Figure 3B,C**). Thus, the pCT provides the core determinants of PKA-C binding, while other regions of SMO may boost the efficiency of the interaction.

To verify the conclusions of our BRET studies biochemically, we tested whether SMO and PKA-C copurify from detergent-solubilized HEK293 cells expressing both proteins. We found that PKA-C copurified with SMO, and the amount is dramatically enhanced by dithiobis(succinimidyl propionate) (DSP), a membrane-permeable amine-specific crosslinker added prior to cell lysis to stabilize protein complexes. In contrast, Nbβ2AR80 did not copurify with SMO **(Figure 3E and Figure 3—figure supplement 2)**. Consistent with our BRET studies, SMO / PKA-C copurification required the SMO pCT **(Figure 3E)**. These data biochemically confirm our BRET findings that SMO and PKA-C interact specifically. SMO / PKA-C complexes may contain additional proteins or lipids. However, SMO and PKA-C copurify in similar quantities, and no other proteins were present at comparable levels other than nonspecific contaminants **(Figure 3E)**, suggesting that SMO and PKA-C interact directly.

### SMO interacts with free PKA-C subunits rather than PKA holoenzymes

Many GPCRs employ A-kinase anchoring proteins (AKAPs) to bind PKA holoenzymes via their PKA-R subunits (Scott and Pawson, 2009; Torres-Quesada et al., 2017). However, our biochemical studies **(Figure 3E)** suggest that SMO interacts directly with free PKA-C without participation from PKA-R. To test this hypothesis, we performed BRET experiments in cells expressing SMO and PKA-C or PKA-R or both **(Figure 4A)**. YFP-tagged PKA-R exhibited only minimal BRET with SMO **(Figure 4B)**, and coexpression of untagged PKA-C did not increase this signal **(Figure 4C, Vehicle)**. These data indicate that PKA-C neither interacts with SMO via PKA-R **(Figure 4A, *iii*)**, nor recruits PKA-R-containing holoenzymes to SMO **(Figure 4A, *ii*)**. SMO therefore does not bind holoenzymes that require cAMP for activation; instead, SMO binds free, catalytically active PKA-C subunits **(Figure 4A, *i*)**.

**Figure 4:**
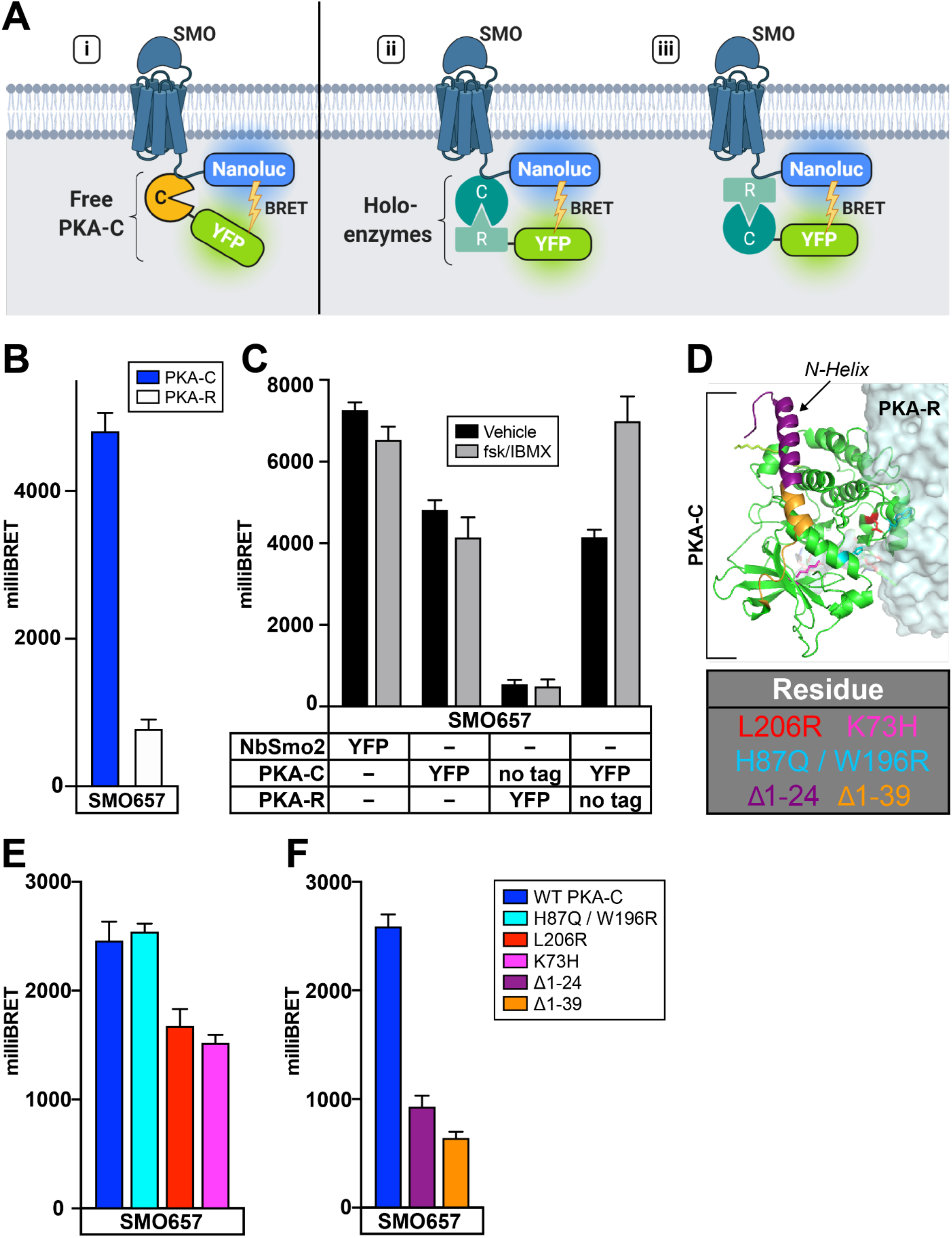
SMO interacts with free PKA-C subunits rather than PKA holoenzymes. (**A**) Schematic of BRET assays to test whether SMO interacts with (i) free PKA-C, or intact PKA holoenzymes via (ii) PKA-C or (iii) PKA-R subunits. (**B**) BRET between a SMO657-nanoluc donor and YFP-tagged PKA-C or PKA-R in HEK293 cells. (**C**) HEK293 cells were transfected with a SMO657-nanoluc donor and untagged or YFP-tagged PKA-C or PKA-R subunits, as described in the table. To stimulate cAMP production, cells were treated for four hours with forskolin (10 μM) + the phosphodiesterase inhibitor isobutylmethylxanthine (IBMX, 1 mM), which blocks cAMP degradation, prior to BRET measurements. (**D**) Structure of PKA holoenzyme (PDB: 4X6R). Key PKA-C residues are colored in the structure and indicated in the table (below). (**E**) BRET between SMO and PKA-C harboring mutations in various regions of the PKA-R binding interface (H87Q/W196R or L206R) or the active site (K73H). (**F**) BRET between SMO and PKA-C harboring deletions of the first 24, or all 39, amino acids from the N-tail. Data are reported as milliBRET ratios and background-subtracted as in Figure 3 (n = 3-6 biological replicates per condition. Error bars = s.e.m.) See Supplemental Table 1 for statistical analysis.

Although SMO / PKA-C complexes do not directly involve PKA-R, cAMP signals may still affect SMO / PKA-C interactions by dissociating holoenzymes, thereby increasing concentrations of free PKA-C within the cell. We therefore examined the effect of cAMP production on SMO / PKA-C BRET in the presence of untagged PKA-R. cAMP production increased BRET under these conditions **(Figure 4C)**. Effects of cAMP production on SMO / PKA-C BRET required PKA-R expression, as expected (**Figure 4C)**. We conclude that cAMP can act on PKA holoenzymes to tune the SMO / PKA-C interaction.

PKA-C forms a bi-lobed structure with an active site involved in substrate binding and phosphoryl transfer on one face and an extended PKA-R binding surface on the opposite face (Taylor et al., 2012, 2013). In addition, an N-terminal tail (“N-tail”) region undergoes myristoylation and mediates interactions with accessory factors **(Figure 4D)** (Bastidas et al., 2012; Pepperkok et al., 2000; Sastri et al., 2005; Tholey et al., 2001). A K73H mutation in the active site (Knighton et al., 1991; Zhang et al., 2015b) modestly inhibited BRET between SMO and PKA-C. An L206R mutation that affects substrate recognition and the PKA-R binding interface (Hannawacker et al., 2019) gave similar results, while H87Q W196R (Orellana and McKnight, 1992), which affects a distinct region of the PKA-R binding interface (Zhang et al., 2015b), did not block BRET **(Figure 4E)**. In contrast, deleting portions of the N-tail significantly reduced BRET between SMO and PKA-C **(Figure 4F)**. These experiments highlight a critical role for the N-tail of PKA-C in mediating interactions with SMO.

### SMO / PKA-C interactions depend on SMO and GRK2/3 activity

PKA phosphorylation of GLI in Hh pathway-responsive tissues or CREB in our heterologous cell assays **(Figure 1)** depends on SMO activity state: it is low when SMO is active, and high when SMO is inactive. One way for PKA phosphorylation of these substrates to reflect SMO activity would be for SMO / PKA-C interactions to increase when SMO is active and decrease when SMO is inactive.

We tested this hypothesis by using BRET to examine the impact of small molecule SMO ligands on SMO interactions with either PKA-C or NbSmo2 (which strongly prefers to bind active SMO over inactive SMO **(Figure 2—figure supplement 1A,B)**). In these experiments, we measured interactions over the full dynamic range of SMO activity by comparing effects of a high-efficacy SMO agonist, SAG21k, to those of KAADcyc. SMO BRET with PKA-C or NbSmo2 was high with SAG21k but decreased with KAADcyc **(Figure 5A and Figure 5—figure supplement 1D)**. SMO modulators produced similar effects on PKA-C or NbSmo2 membrane colocalization **(Figure 5B,C and Figure 5—figure supplement 1A-C)**. In these experiments, SMO modulators exerted somewhat weaker effects on PKA-C than they did on the NbSmo2 positive control. Nevertheless, these effects may be highly relevant under physiological conditions in the presence of other regulatory influences (see “Discussion”). In any event, these data indicate that SMO / PKA-C interactions vary with SMO activity state – they are significantly enhanced when SMO is active.

**Figure 5:**
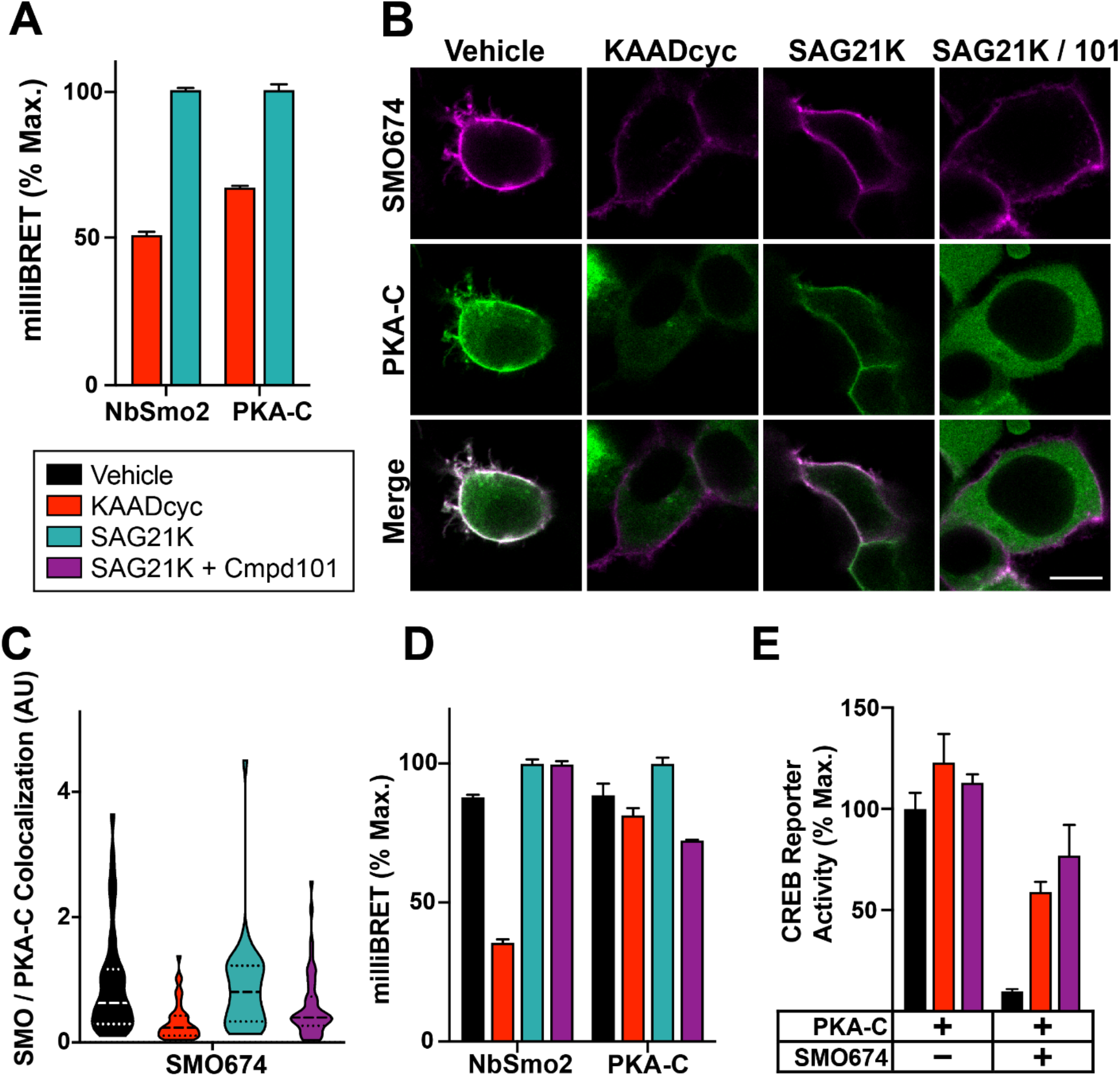
SMO / PKA-C interactions depend on SMO and GRK2/3 activity. (**A**) HEK293 cells transfected with SMO657-nanoluc and PKA-C-YFP were treated with SMO antagonist KAADcyc (1 μM) or agonist SAG21K (1 μM) for one hour prior to BRET measurements. (**B**) Images of HEK293 cells transfected with FLAG-tagged SMO674 (magenta) and YFP-tagged PKA-C (green), and treated with vehicle, KAADcyc (300 nM), or SAG21K (100 nM) alone or with the GRK2/3 inhibitor Cmpd101 (30 μM). Scale bar = 10 μm. (**C**) Quantification of colocalization between SMO and PKA-C for the experiment in (B) (see “Methods’). (**D**) HEK293 cells were transfected with SMO657-nanoluc and YFP-tagged versions of either NbSmo2 or PKA-C, and treated with vehicle, KAADcyc (1 μM), or SAG21K (1 μM) alone or with Cmpd101 (30 μM) for four hours. (**E**) Effect of KAADcyc (1 μM) or Cmpd101 (30 μM) on SMO inhibition of the CREB reporter in HEK293 cells. For (A), and (D), BRET was normalized to 100%, which represents the maximum BRET signal from each set of cells treated with SAG21k. For (E), CREB reporter was normalized to 100%, which reflects reporter activation from PKA-C-transfected cells treated with vehicle. Data in (A), (D), (E): n = 3-6 biological replicates per condition. Error bars = s.e.m.). Data in (C): N = 119-216 cells per condition pooled from two or more independent experiments. See Supplemental Table 1 for statistical analysis.

In considering how activation of SMO leads to enhanced PKA-C binding, GRK family kinases emerged as leading candidates for controlling this process. GRKs selectively phosphorylate the active states of many GPCRs on their intracellular domains, thereby triggering interactions with cytoplasmic regulatory factors (Evron et al., 2012; Homan and Tesmer, 2014; Komolov and Benovic, 2018). In keeping with this paradigm, PKA-C binds active SMO via its pCT. Moreover, GRK2 can phosphorylate active SMO (Chen et al., 2004, 2011), and inhibition of GRK2 and GRK3 strongly disrupts Hh signal transduction (Breslow et al., 2018; Meloni et al., 2006; Philipp et al., 2008; Pusapati et al., 2017, 2018; Zhao et al., 2016).

We tested the effect of GRK2/3 activity on SMO / PKA-C colocalization and binding. The selective GRK2/3 inhibitor Takeda Compound 101 (Cmpd101) inhibited PKA-C colocalization (**Figure 5B, C, and Figure 5—figure supplement 1B**) and BRET **(Figure 5D and Figure 5—figure supplement 1E)** with SMO. As a control, Cmpd101 did not affect BRET with NbSmo2 (**Figure 5D and Figure 5—figure supplement 1E)**. Finally, Cmpd101, like KAADcyc, reversed SMO-dependent inhibition of the CREB reporter (**Figure 5E)**. These findings show that GRK phosphorylation mediates the activity-dependent binding of SMO to PKA-C.

### GRK2/3 phosphorylation of conserved SMO pCT residues mediates PKA-C binding

A parsimonious interpretation of the Cmpd101 results is that PKA-C recruitment to SMO is dependent on GRK2/3-mediated phosphorylation of the SMO intracellular domains. To test this hypothesis, we identified GRK2/3 phosphorylation sites in SMO purified from HEK293 cells stimulated with SAG21k, KAADcyc, or Cmpd101. We then tested whether alanine substitution of the corresponding residues blocked PKA-C interactions.

Phosphoprotein staining of purified SMO samples revealed SMO activity- and GRK2/3-dependent phosphorylation that required the pCT **(Figure 6A)**. Quantitative mass spectrometry (MS) identified seven sites within three clusters (*a:* S560, *b:* S594 / T597 / S599, and *c:* S642 / T644 / T648) exhibiting phosphorylation that depended on SMO and GRK2/3 activity, along with two constitutive phosphorylation sites (S578, S666) **(Figure 6B,C and Figure 6—figure supplement 1A,B)**. Several of these sites are evolutionarily conserved (Maier et al., 2014), particularly in vertebrates **(Figure 6B)**. Four of these sites (S560, S578, T644, and T648) were not detected in previous studies of vertebrate SMO phosphorylation, which involved *in vitro* kinase assays with soluble SMO CT fragments (Chen et al., 2011). In addition, we did not observe phosphorylation at four of the previously mapped sites (S592, S615, S619, S620) (Chen et al., 2011), despite efficient MS coverage of all SMO intracellular regions (data not shown). No SMO activity-dependent GRK2/3-independent phosphorylation events, or vice versa, were detected, suggesting that GRK2/3 are the principal kinases that recognize and phosphorylate active SMO in our experiments.

**Figure 6:**
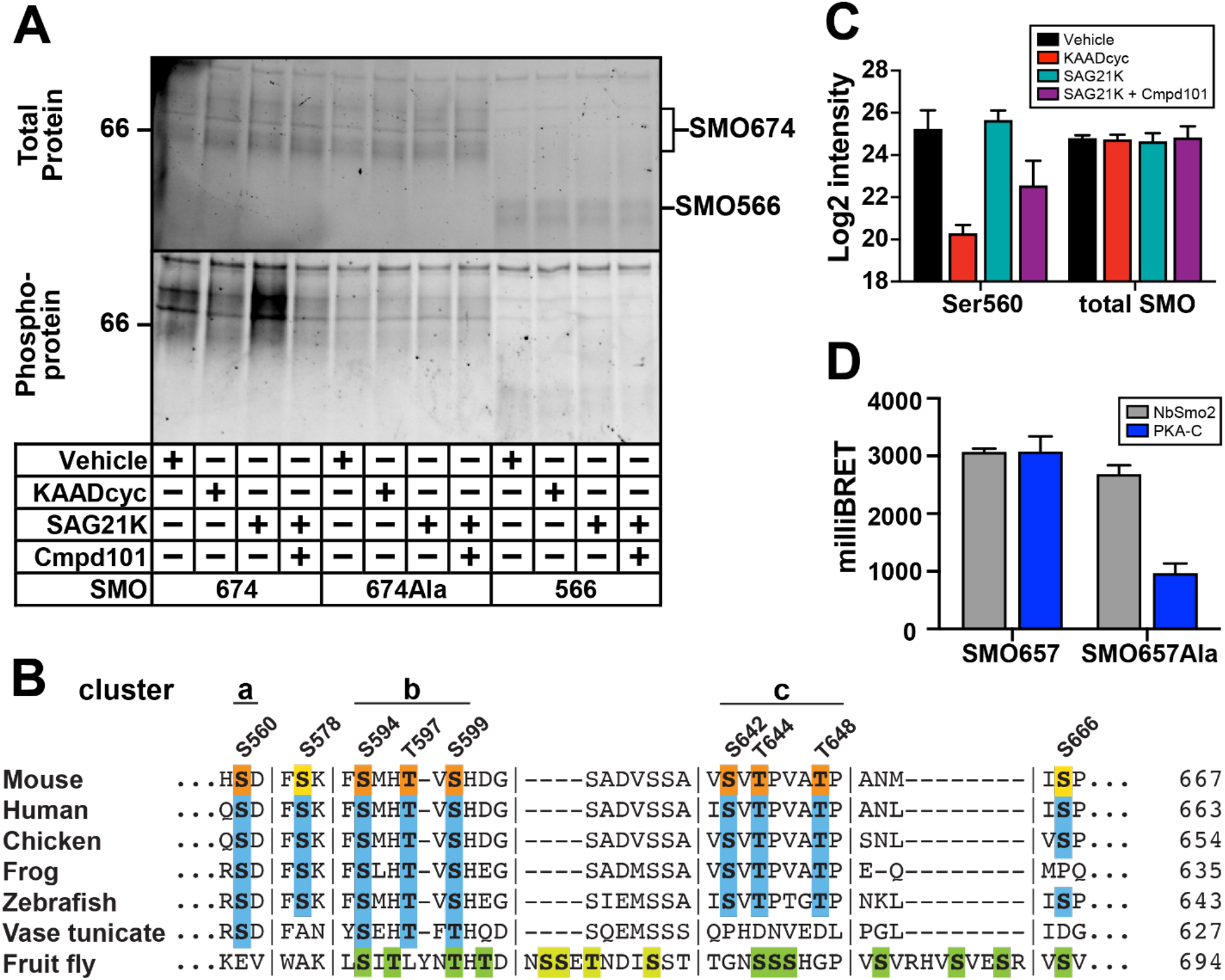
GRK2/3 phosphorylation of conserved SMO pCT residues mediates PKA-C binding. (**A**) HEK293 cells expressing GRK2 and either SMO674 (lanes 1-4), SMO674Ala, which carries mutations in seven GRK2/3 phosphorylation sites (lanes 5-8), or SMO566 (lanes 9-12). Following treatment with SMO modulators or Cmpd101, SMO was isolated via FLAG affinity chromatography and total protein or phosphoprotein was visualized using Stain-Free imaging or Pro-Q Diamond staining, respectively. Although GRKs often phosphorylate GPCRs on the intracellular loops of their 7TM domains (Homan and Tesmer, 2014; Komolov and Benovic, 2018), we did not observe phosphorylation within this region of SMO via phosphoprotein staining (A) or MS (data not shown). Molecular masses are in kDa. (**B**) Clusters of phosphorylated residues identified by MS are labeled above the sequence of mouse SMO. Orange indicates phosphorylation that depends on SMO and GRK2/3 activity, while yellow indicates constitutive phosphorylation. Alignment with SMO from other species reveals sequence conservation (blue), particularly among vertebrates. Green indicates GRK phosphorylation sites previously mapped in Drosophila Smo (ref). Vertical lines indicate breaks in sequence. See Figure 6—figure supplement 1A for complete alignment. (**C**) MS-based quantification of: Left: phosphorylation at S560, one of the activity- and GRK2/3-dependent sites in the SMO pCT. Right: total SMO protein in each sample. “Log2 intensity” is a measurement of the abundance of phosphosites (left) or total protein (right), derived from model-based estimation in MSstats which combines individual peptide intensities (see “Methods”). (**D**) BRET between PKA-C and wild-type SMO674 or SMO674Ala. Data in (C): n = 3 biological and 3 technical replicates per condition. Data in (D): n = 3-6 biological replicates per condition. Error bars = s.e.m.). See Supplemental Table 1 for statistical analysis.

Mutation of the seven SMO activity- and GRK2/3-dependent phosphorylation sites to alanine substantially reduced SMO phosphoprotein staining **(Figure 6A)** and SMO BRET with PKA-C **(Figure 6D)**. As a control, BRET between SMO and NbSmo2 occurred at nearly wild-type levels **(Figure 6D)**. These data indicate that GRK2/3 phosphorylation sites in the pCT domain are critical for PKA-C interaction.

### Hh signal transduction is blocked when SMO cannot bind PKA-C

Our heterologous cell model enabled identification and mapping of a GRK2/3-dependent SMO / PKA-C interaction that interferes with PKA phosphorylation of a heterologous soluble transcription factor. To address whether this mechanism contributes to GLI activation in the Hh pathway, we turned to two models of Hh signal transduction: *i)* activation of a GLI transcriptional reporter in cultured fibroblasts upon treatment with Hh ligands, and *ii)* specification of slow muscle cell subtypes during zebrafish development, which is exquisitely sensitive to Hh pathway activity (Eeden et al., 1996; Stickney et al., 2000; Wolff et al., 2003). Transduction of Hh signals in these models strictly requires SMO, PKA, and GLI (Karlstrom et al., 1999; Lipinski et al., 2008; Taipale et al., 2000; Tuson et al., 2011; Varjosalo et al., 2006; Wolff et al., 2003), and also depends strongly on the presence of primary cilia (Haycraft et al., 2005; Huang and Schier, 2009; Kim et al., 2010; Ocbina and Anderson, 2008).

First, we deleted a small stretch of sequence (SMOΔ570-581) that lies within a region of the pCT critical for interaction with PKA-C **(Figure 3C)**. This SMO deletion abolishes activation of GLI in cultured *Smo^-/-^* fibroblasts (Kim et al., 2009; Varjosalo et al., 2006) **(Figure 7A)** without affecting SMO ciliary localization (Kim et al., 2009). Accordingly, the Δ570-581 deletion severely reduced SMO BRET with PKA-C **(Figure 7B)**. BRET with NbSmo2 was substantially less affected **(Figure 7B)**, suggesting that the defect in PKA-C interaction does not stem from issues with expression or ability to assume an active conformation. Thus, SMOΔ570-581 fails to bind PKA-C and to activate GLI.

**Figure 7:**
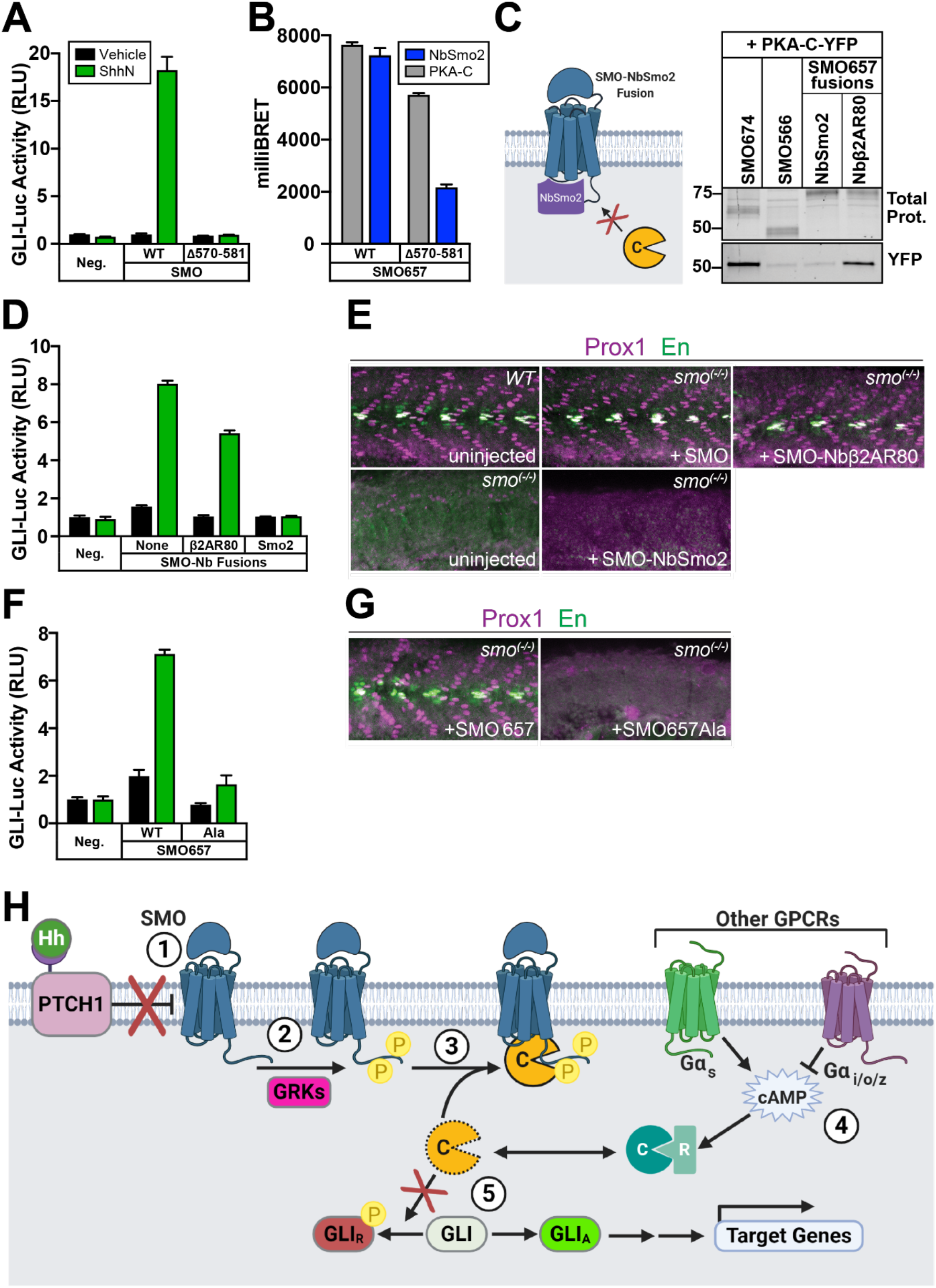
Hh signal transduction is blocked when SMO cannot bind PKA-C. (**A**) GLI transcriptional reporter assay in Smo^-/-^ mouse embryonic fibroblasts (MEFs) expressing wild-type SMO or SMOΔ570-581, treated with control or Sonic hedgehog (ShhN) conditioned medium. GFP serves as a negative control (“Neg.”). RLU = relative luciferase units. (**B**) BRET in HEK293 cells between nanoluc-tagged wild-type or Δ570-581 forms of SMO657 as donor, with YFP-tagged NbSmo2 or PKA-C as acceptor. (**C**) Left: Schematic of SMO-NbSmo2 fusion, predicted to block interactions with SMO that require the intracellular face of the 7TM domain. Right: YFP-tagged PKA-C was coexpressed in HEK293 cells with FLAG-tagged SMO674 (lane 1), SMO566 (lane 2), SMO657-NbSmo2 (lane 3), or SMO657-Nbβ2AR80 (lane 4). Following cell lysis and FLAG purification of SMO complexes, samples were separated on SDS-PAGE. Fluorescence scans of total protein (top) and YFP (bottom) in FLAG eluates are shown. Note that DSP crosslinker was not used in this experiment; thus, copurification of PKA-C was less efficient than in Figure 2F. Molecular masses are in kDa. (**D**) GLI transcriptional reporter assay in Smo^-/-^ MEFs expressing fusions to NbSmo2 or Nbβ2AR80. Non-Nb-fused SMO (“none”) serves as a positive control. (**E**) Confocal images of whole-mount wild-type zebrafish embryos or smo^-/-^ mutants injected with mRNAs encoding either wild type SMO, SMO-Nbβ2AR80, or SMO-NbSMO2. Embryos 26 hours post-fertilization were fixed and stained with antibodies against Prox1 (magenta) or Engrailed (En, green) to mark populations of muscle fiber nuclei. (**F**) GLI transcriptional reporter assay in Smo^-/-^ MEFs expressing wild-type SMO657 (“WT”) or SMO657Ala. (**G**) Zebrafish were injected with mRNAs encoding wild-type SMO657 or SMO657Ala, then stained for muscle fiber nuclei as described in (E). (**H**) Proposed model for SMO activation of GLI via PKA-C membrane sequestration: (1) Hh proteins bind to and inhibit PTCH1, inducing an activating conformational change in SMO; (2) Active SMO is recognized and phosphorylated by GRKs; (3) Phosphorylated SMO recruits PKA-C to the membrane, preventing PKA-C from phosphorylating and inhibiting GLI; (4) GPCRs that couple to Gα_s_ (such as GPR161) or Gα_i/o/z_ (perhaps including SMO itself) can raise or lower cAMP levels, respectively, thereby affecting SMO / PKA-C interactions by regulating the size of the free PKA-C pool. (5) GLI is converted from a repressed (GLI_R_) to an active (GLI_A_) form and regulates transcription of Hh target genes. Data in (A), (B),(D), (F): n = 3 biological replicates per condition. Error bars = s.e.m. Data in (E), (G): n = 78 (SMO), 61 (SMO-NbSmo2), 63 (SMO-Nbβ2AR80), 62 (SMO657), 70 (SMO657Ala), 100 (uninjected). See Supplemental Table 1 for statistical analysis.

Next, we harnessed our insight that SMO / PKA-C interactions depend on SMO activity and GRK2/3 phosphorylation to design a different non-PKA-C-binding SMO mutant. The intracellular region of the SMO 7TM domain changes conformation dramatically upon SMO activation (Deshpande et al., 2019). This same region is also necessary for recruitment of GRKs to the active states of other GPCRs (Homan and Tesmer, 2014; Komolov et al., 2017). To assess whether SMO interactions with PKA-C also required this region, we fused NbSmo2 to the end of the CT. As a result, NbSmo2 is expected to bind SMO and dissociate minimally, if at all; interactions with the 7TM domain’s intracellular region that involve other proteins, domains of SMO, or both, will be efficiently blocked **(Figure 7C, left)**. SMO-NbSmo2 failed to bind PKA-C, while a negative control fusion of similar size and expression level, SMO-Nbβ2AR80, bound robustly **(Figure 7C, right)**. In cultured *Smo^-/-^* fibroblasts, SMO-NbSmo2 also failed to stimulate GLI-dependent transcription in response to Hh ligands, whereas SMO-Nbβ2AR80 still mediated strong transcriptional responses **(Figure 7D)**. In zebrafish, expression of mRNA encoding wild-type SMO or SMO-Nbβ2AR80 restored Hh pathway-dependent muscle development to *smo^-/-^* embryos, whereas SMO-NbSmo2 did not **(Figure 7E)**. Control experiments confirmed that SMO-NbSmo2 and SMO-Nbβ2AR80 accumulate normally in cilia in response to SMO activation (data not shown). These findings argue that blockade of Nb-binding regions in the SMO 7TM domain hinders interactions with PKA-C and activation of GLI. They also further establish a correlation between SMO / PKA-C binding and GLI activation.

Lastly, we mutated the seven critical GRK2/3-dependent phosphorylation sites in SMO. These mutations not only dramatically impaired GLI transcription in cultured *Smo^-/-^* fibroblasts **(Figure 7F)**, but also failed to restore normal muscle specification to *smo^-/-^* zebrafish embryos **(Figure 7G)**.

Taken together, these results highlight an important role for GRK2/3-mediated SMO / PKA-C binding and subsequent PKA-C membrane sequestration in controlling GLI activation in cellular models as well as embryonic patterning *in vivo*.

## DISCUSSION

We have identified and characterized a novel interaction between vertebrate SMO and PKA-C, demonstrated how this recruits PKA-C to membranes, and showed how it can dramatically affect the activity of PKA-regulated transcription factors as well as GLI-dependent outputs in cultured cells and *in vivo*. These insights enable a deeper understanding of how Hh signal transduction orchestrates cell proliferation and differentiation events in nearly all of its biological roles.

Based on these findings, we propose the following mechanism **(Figure 7H)**. In the pathway “off” state, SMO is inactive and inefficiently binds PKA-C. PKA-C is thus available to phosphorylate and inactivate GLI. In the pathway “on” state, SMO activation triggers GRK phosphorylation of the pCT, increasing PKA-C binding and siphoning PKA-C to the membrane. SMO-bound PKA-C cannot access soluble GLI substrates. This leads to loss of inhibitory GLI phosphorylation, which could occur via GLI protein turnover (Hui and Angers, 2011; Humke et al., 2010; Niewiadomski et al., 2013), tonic action of phosphatases on GLI (Zhao et al., 2017), or both.

Our proposed mechanism helps to explain key aspects of SMO-to-GLI communication not easily reconciled with existing models. Prior studies have invoked a variety of explanations for how SMO might activate GLI (Kong et al., 2019), including: *i)* utilizing canonical Gα_i/o/z_ and cAMP-dependent pathways to inhibit PKA (Ayers and Thérond, 2010); *ii)* controlling ciliary cAMP, and thus PKA-C, by influencing the ciliary localization of GPR161, a constitutively active GPCR (Mukhopadhyay et al., 2013); *iii)* interacting with βarrestin1/2 (Kovacs et al., 2008) or Ellis-van Creveld protein 2 (EVC2) (Dorn et al., 2012), which might regulate GLI by as-yet-undefined mechanisms. However, pharmacological or genetic inactivation of Gα_i/o/z_ signaling does not prevent SMO from activating GLI (Low et al., 2008; Regard et al., 2013; Riobo et al., 2006). In addition, whether SMO activation reduces ciliary cAMP remains controversial (Jiang et al., 2019; Moore et al., 2016; Tschaikner et al., 2020). Finally, mouse knockouts of *βarrestin1/2* (Zhang et al., 2010), *EVC2* (Zhang et al., 2015a), or *GPR161* (Hwang et al., 2018; Mukhopadhyay et al., 2013; Shimada et al., 2018), fail to exhibit the severe, widespread developmental defects expected with disruption of core Hh pathway components (Goodrich et al., 1996; Tuson et al., 2011; Zhang et al., 2001). Therefore, existing models neither fully explain how SMO activates GLI, nor rule PKA in or out as a mediator of this process. Our work reveals a route by which SMO can affect PKA substrate phosphorylation that does not require any of these factors, offering an explanation for prior conflicting observations.

### GRKs and Hh signal transduction

Our work also sheds light on how GRKs control Hh signal transduction. Pathway activation strongly requires these kinases in vertebrates, but their underlying target and mechanism of action remained poorly defined (Kong et al., 2019; Meloni et al., 2006; Pusapati et al., 2018; Sharpe and de Sauvage, 2018; Zhao et al., 2016). While GRKs can phosphorylate SMO (Chen et al., 2004), mutation of the previously mapped GRK sites (Chen et al., 2011) to alanine does not disrupt embryonic patterning *in vivo* (Zhao et al., 2016). Consequently, it was unknown whether the physiological target of GRKs in the Hh pathway is SMO or a different protein altogether (Kong et al., 2019; Pusapati et al., 2018; Sharpe and Sauvage, 2018; Zhao et al., 2016). Furthermore, how GRK phosphorylation of its substrate(s) regulates GLI activity remained unclear. A key limitation is that prior studies defined GRK sites based largely on *in vitro* kinase assays with soluble SMO CT fragments (Chen et al., 2011). GRKs lack a strict consensus motif, and capturing physiological activity-induced phosphorylation of GPCRs requires their 7TM domains to be embedded in a membrane lipid environment (DebBurman et al., 1996; Inagaki et al., 2012, 2015; Komolov et al., 2017), making phenotypic interpretation of existing alanine mutants difficult. Here, we studied phosphorylation of SMO (with an intact 7TM domain) expressed in mammalian cells. We also used specific pharmacological agents to define the activity- and GRK-dependence of SMO phosphorylation events. Our analysis revealed several GRK phosphorylation sites in the pCT that were not previously reported. Mutation of the sites we identified strongly affects GLI activation in cultured cells and *in vivo*, indicating that GRK phosphorylation of SMO is in fact critical for Hh signal transduction. Although GRKs may play multiple roles in Hh signal transduction (Pusapati et al., 2018; Sharpe and De Sauvage, 2018; Zhao et al., 2016), our PKA-C sequestration model is particularly appealing because it directly links GRK phosphorylation of SMO to stimulation of GLI.

### Structural determinants of the SMO / PKA-C complex

SMO activation of GLI requires the pCT domain (Kim et al., 2015; Varjosalo et al., 2006), but for reasons that have remained incompletely understood. The pCT contains essential ciliary trafficking motifs (Kim et al., 2015). However, pCT mutations such as Δ570-581 disrupt GLI activation without affecting SMO ciliary trafficking (Kim et al., 2009), indicating that the pCT plays additional indispensable roles in GLI activation besides controlling SMO ciliary localization. Here we show that one such function is to bind and sequester PKA-C when SMO and GRK2/3 are active. Structures of the pCT or of the SMO / PKA-C complex have not yet been reported. However, our mutational analysis suggests that this complex involves, at minimum, the phosphorylated pCT of SMO and the N-tail domain of PKA-C. Membrane lipid interactions may also contribute to these complexes, as the N-tail is myristoylated which can increase PKA-C membrane association in some settings (Bastidas et al., 2012; Gaffarogullari et al., 2011; Tillo et al., 2017; Zhang et al., 2015b). Intriguingly, recent structures of non-SMO GPCRs in complex with βarrestins have also revealed critical interactions with the receptor’s phosphorylated cytoplasmic tail and lipids in the surrounding membrane (Huang et al., 2020; Staus et al., 2020). Thus, distinct GPCRs may engage a diverse set of downstream effectors, such as βarrestins or PKA-C, using similar structural principles.

### SMO control of PKA localization is an evolutionarily conserved rheostat

The Hh pathway controls development and regeneration throughout the animal kingdom (Ingham et al., 2011), but whether the underlying transduction mechanism is conserved remains a matter of debate (Huangfu and Anderson, 2006). Recent studies of SMO communication with GLI have emphasized aspects of the Hh pathway that are uniquely important to mammals but not insects, such as the primary cilium (Gigante and Caspary, 2020; Goetz and Anderson, 2010; Huangfu and Anderson, 2006). In contrast, the SMO / PKA-C interaction we describe here is conserved in *Drosophila* (Li et al., 2014; Ranieri et al., 2014). This interaction promotes *Drosophila* Hh pathway activation in part by titrating PKA-C out of a protein complex that promotes phosphorylation and inhibition of the GLI ortholog Ci (Li et al., 2014; Ranieri et al., 2014). This action is strikingly parallel to effects of SMO / PKA-C interactions on CREB reporter activation and GLI-dependent transcription observed here. Thus, SMO may utilize PKA to communicate with GLI via mechanisms that are more similar between species than previously appreciated. One evolutionary advantage to a mechanism based on direct SMO / PKA-C interactions is that it ensures graded PKA inhibition over a broad range of SMO activity levels. As a result, SMO can accurately translate differences in amounts of extracellular Hh proteins (via PTCH1 binding and inactivation) into discrete changes in GLI activity. This is essential for Hh proteins to function in gradients as concentration-dependent (morphogenetic) signals in the limb bud, neural tube, and elsewhere. In contrast, with cascades that involve intermediary components present in limiting amounts, downstream responses may reach maximal levels even when upstream receptors are not fully activated (Kenakin, 2008), causing a loss of signal fidelity at high levels of receptor activity.

### The role of the cilium in SMO regulation of PKA activity

In our HEK293 model, SMO activation triggered changes in the interactions, localization, and activity of a substantial fraction of cellular PKA-C. In contrast, under physiological conditions, SMO would mainly act on the much smaller pool of PKA-C in the primary cilium. In this manner, SMO could specifically regulate GLI transcription factors without affecting PKA-dependent processes elsewhere in the cell (Jiang et al., 2019; Mukhopadhyay et al., 2013; Tschaikner et al., 2020; Tuson et al., 2011). Upon Hh pathway activation, SMO not only changes conformation within the cilium (Rohatgi et al., 2009; Wilson et al., 2009), but also accumulates to high levels in the ciliary membrane (Corbit et al., 2005; Kim et al., 2009; Rohatgi et al., 2007). This increase in SMO abundance, along with the SMO activity-dependent binding events described in our study, may synergize to effectively sequester ciliary PKA-C at the membrane and away from GLI proteins in the interior of the cilium (cilioplasm). Such a process could also involve transfer of PKA-C out of ciliary protein complexes that facilitate GLI phosphorylation and inhibition, and into SMO-containing complexes that do not. Curiously, SMO and PKA-C do not appear to overlap within the cilium in standard immunofluorescence microscopy; PKA-C is enriched at the basal body of the cilium, while SMO is in the membrane (Barzi et al., 2009; Tuson et al., 2011). Nevertheless, a recent study, using an elegant strategy to specifically inhibit PKA-C at defined locations within the cilium, found evidence for a labile pool of cilioplasmic PKA-C that is critical for GLI regulation (Mick et al., 2015). Along similar lines, PKA-C activity has been detected within the cilioplasm using a FRET-based sensor of PKA substrate phosphorylation (Moore et al, 2016). SMO may activate GLI by sequestering this pool of PKA-C via the mechanism described in our study. In the future, live-cell super-resolution microscopy may enable visualization of cilioplasmic PKA-C and evaluation of its distribution within the cilium before and after SMO activation.

### SMO control of GLI may require several mechanisms acting in concert

The mechanism we identified likely acts together with other processes to enable SMO activation of GLI. For example, SMO or other GPCRs (such as GPR161) may still utilize G proteins to set levels of cAMP, and thus levels of free PKA-C, within a critical range that allows SMO / PKA-C interactions to affect GLI activity (**Fig. 7H)**. Within this range, PKA-C could access GLI when SMO is inactive but undergo efficient membrane sequestration when SMO is active. This is consistent with observations that manipulation of cAMP signals, via expression of dominant negative (non-cAMP-binding) PKA-R constructs or treatment with forskolin (Eeden et al., 1996; Hammerschmidt et al., 1996; Taipale et al., 2000), strongly increases or decreases GLI activity, respectively. Along similar lines, knockout of GPR161 elevates Hh pathway activity in some settings (Hwang et al., 2018; Mukhopadhyay et al., 2013; Pusapati et al., 2018; Shimada et al., 2018). Thus, a number of processes may cooperate with the SMO mechanism described here to create a robust PKA-C activity switch. In this regard, while SMO or GRK2/3 modulators exert modest effects on SMO / PKA-C interactions and colocalization in some of our experiments, they may dramatically affect PKA-C substrate phosphorylation under physiological conditions where other regulatory influences are present.

### GPCR signaling without second messengers

The mechanism we describe here for the Hh pathway may apply more generally to communication between GPCRs and PKA. These receptors and effectors participate in numerous signaling cascades that mediate an extraordinarily diverse range of biological processes (Lefkowitz, 2000, 2002; Scott and Pawson, 2009; Taylor et al., 2013). Yet, it remains unclear how communication between just two types of signaling molecules can produce such a vast array of cellular and physiological outputs. Prior studies have focused largely on indirect modes of GPCR-PKA communication involving G proteins, cAMP, and AKAP adaptors (Lefkowitz, 2002; Scott and Pawson, 2009). In contrast, our study describes an alternative mechanism, based on direct PKA-C interactions with an active GPCR. This mechanism may act in concert with classical second-messenger signals to bias phosphorylation of PKA substrates toward or away from specific subcellular locations. Such receptor-mediated PKA sequestration may constitute a broader theme among GPCRs in the cilium, as was recently shown for GPR161, which encodes an AKAP domain in its intracellular C-terminus that binds and recruits PKA-R subunits to the ciliary membrane (Bachmann et al., 2016). The additional level of spatial regulation gained from these strategies may allow GPCR-containing pathways throughout the cell to encode new types of downstream responses, thereby permitting control of an expanded array of biological outputs.

## ACKNOWLEDGMENTS

We thank A. Inoue for providing HEK293 Ga-null cells, and S. Lusk and K. Kwan for providing *smo^hi1640^* zebrafish. We thank D. Julius, M. He, and S. Nakielny for providing feedback on the manuscript. C.D.A. and I.B.N. acknowledge support from the Undergraduate Research Opportunities Program at the University of Utah. D.K.S. acknowledges support from the NIH Developmental Biology Training Grant at the University of Utah (T32HD007491). I.D. acknowledges support from the Swiss National Science Foundation. N.J.K. and R.H. are supported by the US Department of Defense Advanced Research Projects Agency (HR0011-19-2-0020, HR001119S0092-FP-FP-002). A.M. acknowledges support from the Pew Charitable Trusts. D.J.G. and B.R.M acknowledge support of funds in conjunction with grant P30CA042014 awarded to Huntsman Cancer Institute and to the NC Program at Huntsman Cancer Institute. This work was supported by NIH award 1R35GM133672 (B.R.M.) and an ACS Institutional Research Grant award (B.R.M.).

## AUTHOR CONTRIBUTIONS

B.R.M., C.D.A., J.T.H., A.M., R.H., and D.J.G. conceived and designed the project, B.R.M., A.M., R.H., N.J.K., and D.J.G. provided overall project supervision. C.D.A. and J.T.H. conducted BRET experiments. J.T.H. conducted CREB-based reporter studies. D.S.H. conducted cultured cell imaging experiments. J.Z. conducted SMO / PKA-C copurification experiments. D.K.S. and D.J.G. conducted zebrafish injections and immunohistochemistry. J.X. and R.H. conducted mass spectrometry experiments and data analysis. I.D. designed SMO purification constructs and analyzed SMO-Nb complexes by size exclusion chromatography. A.M. and J.L. purified NbSmo2 and performed FACS-based NbSmo2 binding assays. J.L.C., S.L.S., I.B.N., and B.R.M. performed GLI-luciferase assays. J.L.C., I.B.N., and M.F.W. analyzed imaging data (with guidance from D.S.H.). J.T.H. and I.B.N. assembled manuscript figures, and B.R.M. and C.D.A. wrote the manuscript with input from all authors.

## SUPPLEMENTAL INFORMATION

### Supplemental information on experimental model system

Hh signal transduction is often studied using GLI transcriptional readouts (Taipale et al., 2000). These readouts present two major obstacles for determining whether SMO inhibits PKA to activate GLI. First, GLI transcription is strongly affected by manipulation of either SMO or PKA (Hui and Angers, 2011; Kong et al., 2019; Taipale et al., 2000), complicating efforts to determine whether SMO and PKA reside in the same linear pathway or constitute two separate influences on GLI. Second, during Hh signal transduction, SMO and GLI are subject to elaborate ciliary trafficking mechanisms (Gigante and Caspary, 2020; Goetz and Anderson, 2010) that are incompletely understood and difficult to disentangle from the events occurring immediately downstream of SMO activation. To strip away these potentially confounding factors, we developed a heterologous HEK293 model for SMO regulation of PKA activity. This approach permits simple, direct measurements of SMO effects on PKA, independent of ciliary trafficking or other intermediate steps (Myers et al., 2017). We used CREB transcription to monitor PKA phosphorylation in HEK293 cells. CREB, like GLI, is a soluble transcription factor regulated by PKA phosphorylation (although PKA phosphorylation activates CREB but inhibits GLI.) However, CREB is not known to be subject to the other major mechanisms that regulate GLI activity (Hui and Angers, 2011; Shaywitz and Greenberg, 1999). Therefore, any effects of SMO on CREB transcription would provide evidence that SMO can control PKA. In addition, unlike GLI, activation of CREB transcription factors is not reported to require the primary cilium. As a result, we can directly study SMO effects on PKA function in the absence of ciliogenesis or ciliary protein trafficking processes.

### Statistics

Representative data from at least two independent trials are shown. Data is reported as the mean of at least three biological replicates with error bars representing standard error of the mean. Statistical tests were performed as described in Supplemental Table 1.

## SUPPLEMENTAL FIGURES

**Figure 1—figure supplement 1:**
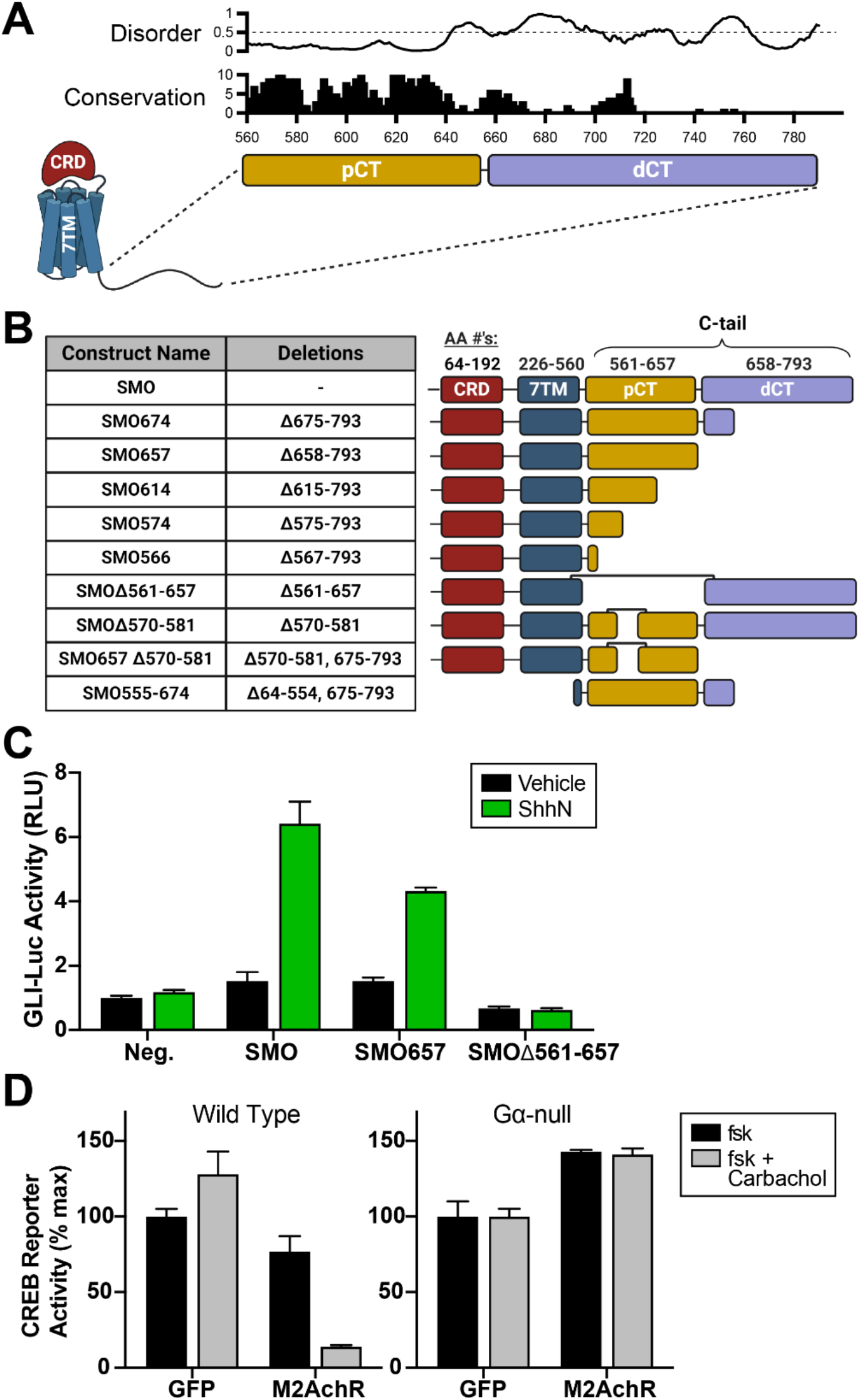
SMO expression constructs and controls for CREB-reporter-based PKA activity assay. (**A**) Jalview conservation score (0-10) and DISOPRED disorder score (0-1, with values >0.5 indicative of disorder) are plotted on the y-axes, while SMO amino acid numbering is plotted on the x-axis. Locations of proximal C-tail (pCT, mustard) and distal C-tail (dCT, lavender) are indicated below the graphs. (**B**) Table of SMO constructs used in this study. Note that CRD (red) and 7TM (blue) domains are not drawn to scale. We used SMO657, truncated immediately after the pCT, for many of our cell-based experiments, based on its ability to recapitulate >70% of the activity of wild-type SMO in GLI reporter assays (see (**C**)). However, secondary structure predictions revealed a conserved region between amino acids 657-674, predicted to be ordered and to correspond to the end of a helix (data not shown). Because these parameters might help to increase the biochemical stability of SMO, we extended the construct boundary (SMO674) in any experiments requiring purification of SMO protein. (**C**) Smo^-/-^ mouse embryonic fibroblasts (MEFs) were transfected with GLI-luciferase reporter plasmid, along with a GFP negative control (“Neg.”), full-length SMO, or truncation mutants lacking the dCT (SMO657) or pCT (SMOΔ561-657). Following transfection, cells were stimulated with conditioned medium containing the N-terminal signaling domain of Sonic hedgehog (ShhN, green) or control, non-ShhN-containing conditioned medium (vehicle, black). RLU = relative luciferase units. (**D**) Wild type (left) or Gα-null (right) HEK293 cells transfected with GFP (negative control) or M2AchR expression plasmids, stimulated with forskolin (black, 500 nM for wild-type cells, 80 μM for Gα-null cells) in the presence or absence of carbachol (gray, 3 μM). Note that substantially less forskolin is needed to induce cAMP signals in wild-type HEK293 cells compared to Gα-null cells due to the presence in the former of Gα_s_, which binds to and sensitizes adenylyl cyclase (AC) to forskolin treatment (A. Inoue, personal communication.) In addition, basal reporter activity in Gα-null cells is higher following M2AchR expression, but the interpretation of this effect is uncertain because it is not altered by treatment with carbachol. Data are normalized to 100%, which reflects reporter activation from PKA-C-transfected cells treated with vehicle (n = 3 biological replicates per condition, error bars represent s.e.m.). See Supplemental Table 1 for statistical analysis.

**Figure 2—figure supplement 1:**
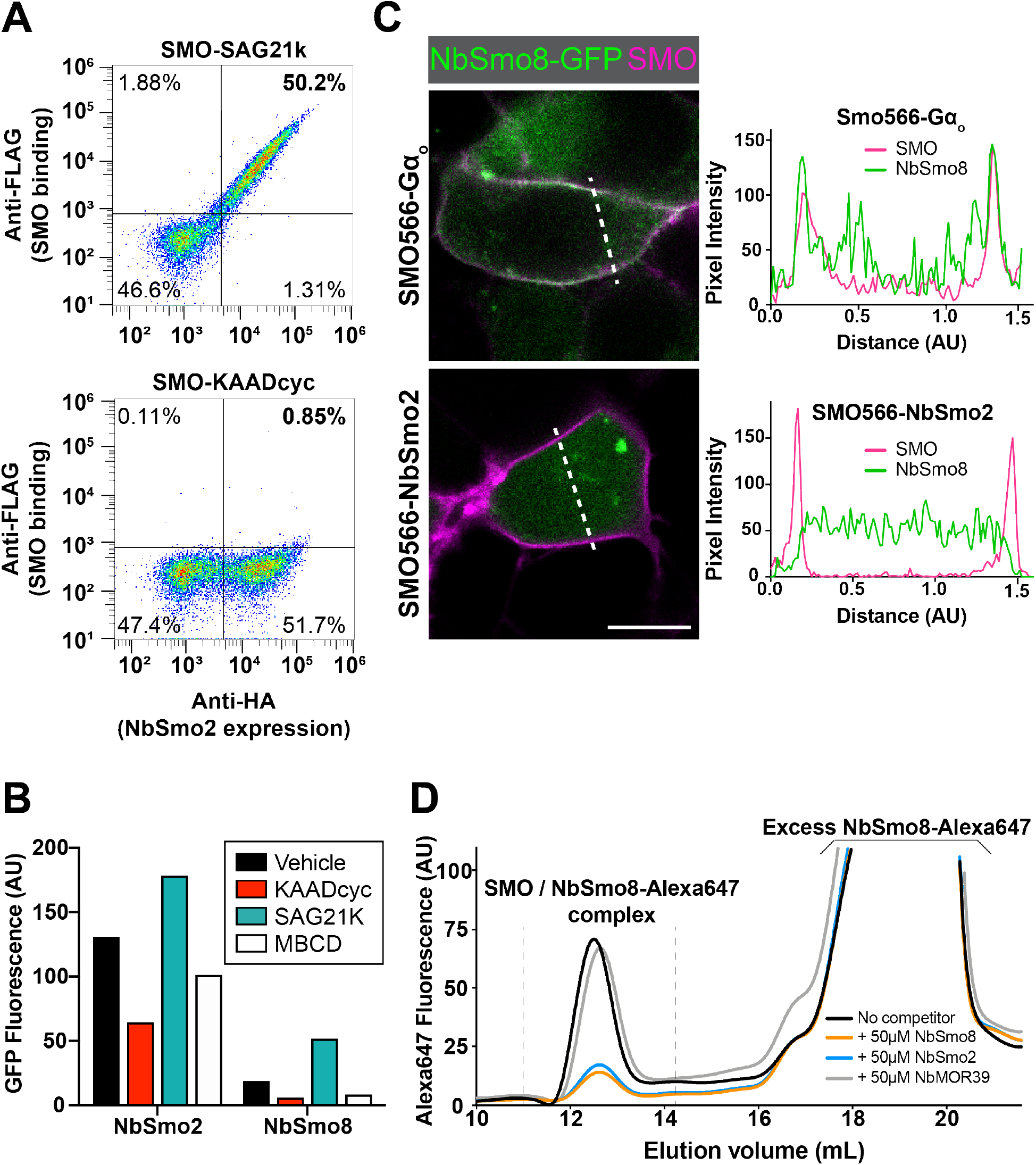
Confocal imaging line scans and Nb2 selection. (**A**) Binding of NbSmo2, displayed on the surface of yeast (Deshpande et al., 2019), to purified, detergent-solubilized SMO-agonist (SAG21k) complexes or SMO-antagonist (KAADcyc) complexes in solution, was assessed by flow cytometry. Note that this experiment used SMO566, which lacks the entire cytoplasmic tail. (**B**) FLAG-tagged SMO566-Gα_o_ was expressed in HEK293 cells alone or with GFP-tagged NbSmo2, NbSmo8, or Nbβ2AR80. Following treatment with SMO agonist (SAG21k, 1 μM), antagonist (KAADcyc, 1 μM), or methyl-beta-cyclodextrin (MBCD, 8 mM, which extracts SMO sterol agonists from membranes (Myers et al., 2013)), SMO-Nb complexes were isolated from detergent-solubilized cells via FLAG affinity chromatography and Nb levels measured via GFP fluorescence quantification. Ratios of GFP fluorescence in FLAG eluates, normalized to GFP fluorescence in each lysate before affinity chromatography, are reported. (**C**) NbSmo8-GFP colocalization with SMO566-NbSmo2 fusion at the cell membrane. The presence of NbSmo2 is predicted to prevent binding of NbSmo8 to SMO if the Nbs bind to overlapping epitopes. SMO566-Gα_o_ serves as a positive control. Line scan analysis is shown to the right of each merged image, with a dotted line indicating the location of the scan. (**D**) In vitro binding of Alexa647-labeled NbSmo8 to SMO566 in the presence of non-fluorescent NbSmo2 competitor, as assessed by fluorescence detection size exclusion chromatography. Non-fluorescent NbSmo8 or NbMOR39 (which binds a non-SMO GPCR (Huang et al., 2015)) serve as positive and negative controls for NbSmo8 competition binding, respectively.

**Figure 2—figure supplement 2:**
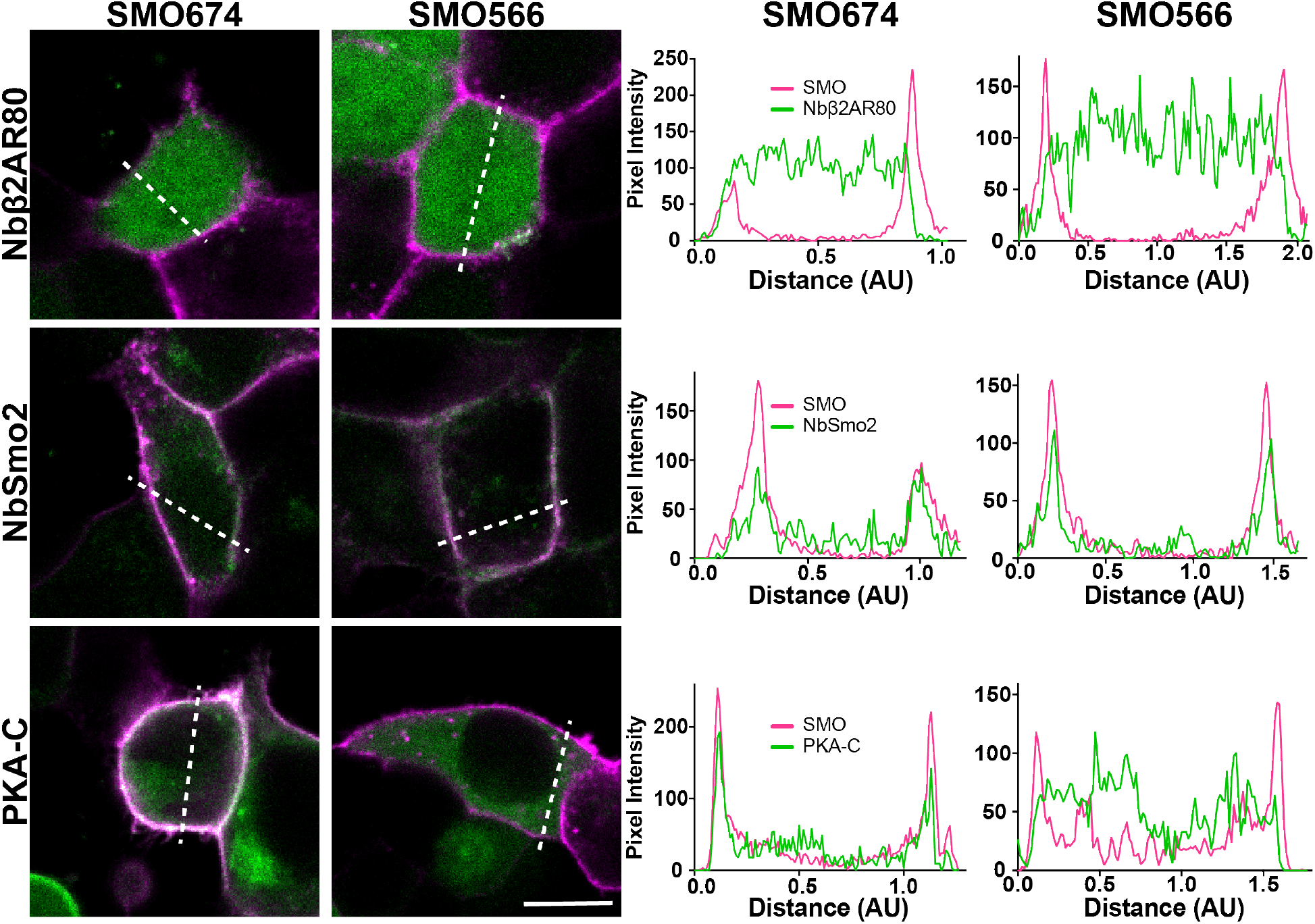
Line scans. Line scans for colocalization images in **Figure 2A** and. **Figure 2B**. Colors are the same as described in the main figure panels. Dotted line indicates location of the scan.

**Figure 3—figure supplement 1:**
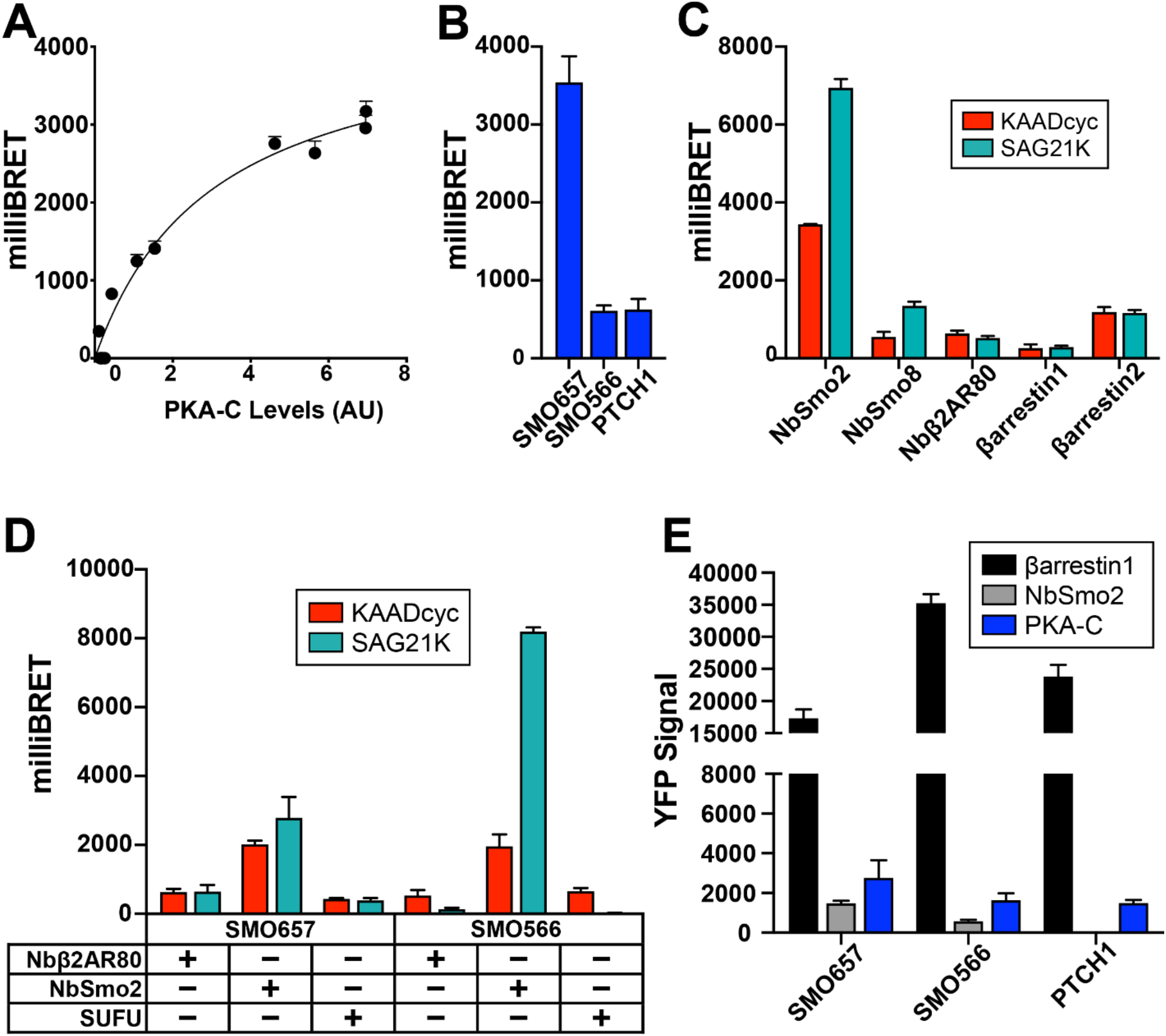
Controls for BRET and SMO / PKA-C BRET studies. (**A**) Saturation analysis of BRET between SMO and PKA-C. A fixed amount of SMO BRET donor was cotransfected with increasing amounts of PKA-C BRET acceptor. The x-axis reflects levels of PKA-C, normalized to levels of SMO as described in Methods (AU = arbitrary units), and the y-axis reflects the BRET ratio (in milliBRET units). (**B**) BRET between YFP-tagged PKA-C and nanoluc-tagged SMO657, SMO566, or PTCH1. (**C**) SMO BRET with NbSmo2, NbSmo8, Nbβ2AR80, βarrestin-1, or βarrestin-2. To determine if BRET depends on SMO activity, cells were treated for 1 hr. with SAG21K (1μM) or KAADcyc (1μM). (**D**) BRET using SMO657 or SMO566 as donors and Nbβ2AR80, NbSmo2, or SUFU as acceptors, performed as described in (C). **(E)** Levels of BRET acceptor for the experiment shown in **Figure 3B**, measured as YFP fluorescence prior to addition of nanoluc substrate, with background subtracted. See Supplemental Table 1 for statistical analysis.

**Figure 3-figure supplement 2:**
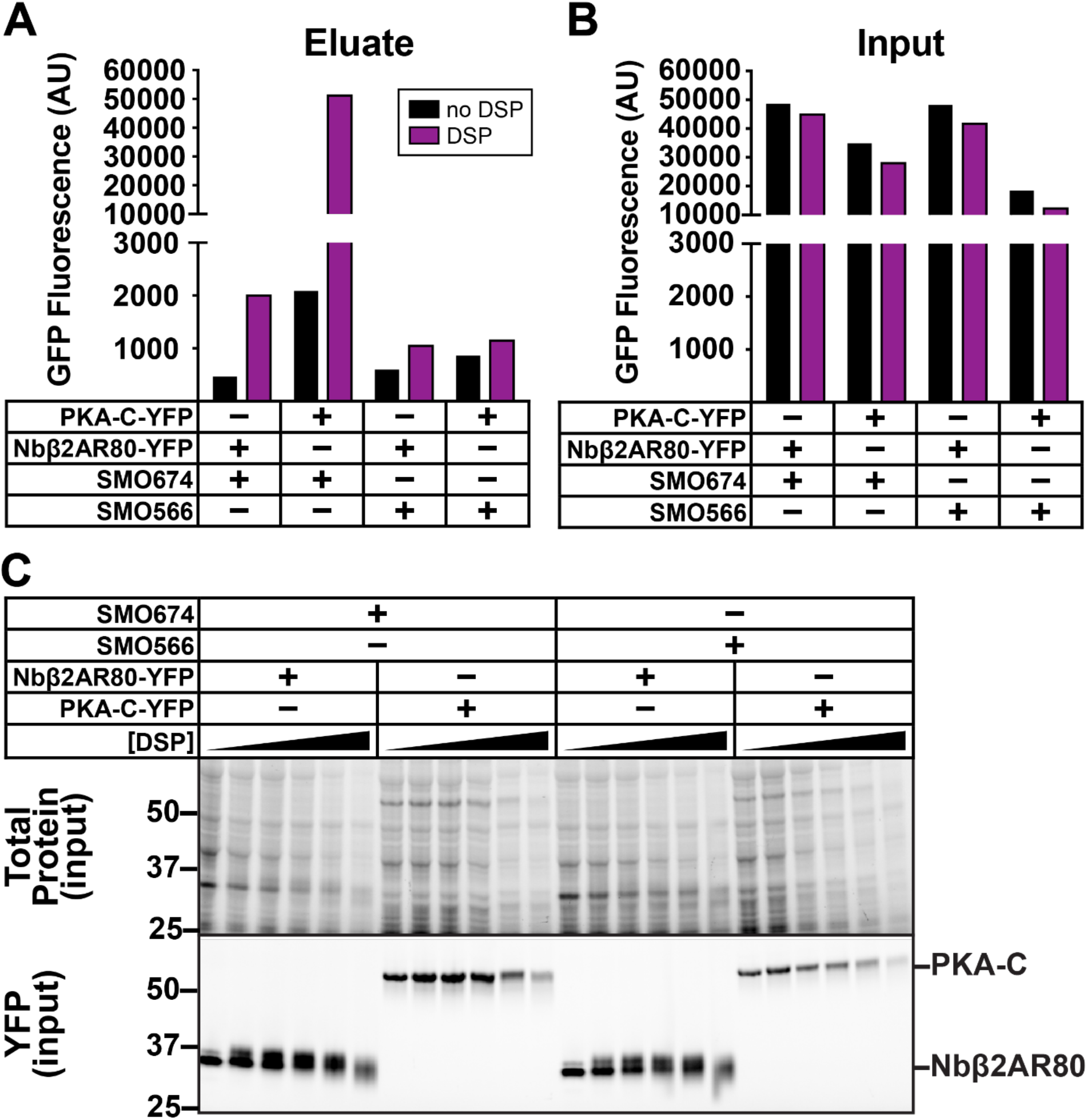
Controls for SMO / PKA-C crosslinking studies. YFP quantification of (**A**) FLAG elution fractions or (**B**) input fractions from SMO / PKA-C copurification performed in the absence (black) or presence (purple) of 0.5 mM DSP crosslinker, as presented in **Figure 3E**. (**C**) Protein gels of input fractions for the experiment shown in **Figure 3E**. The decrease in soluble protein yields in total cell lysates that occurs at high DSP concentrations may be due to crosslinker-induced protein aggregation and loss of solubility. Molecular masses are in kDa.

**Figure 5-figure supplement 1:**
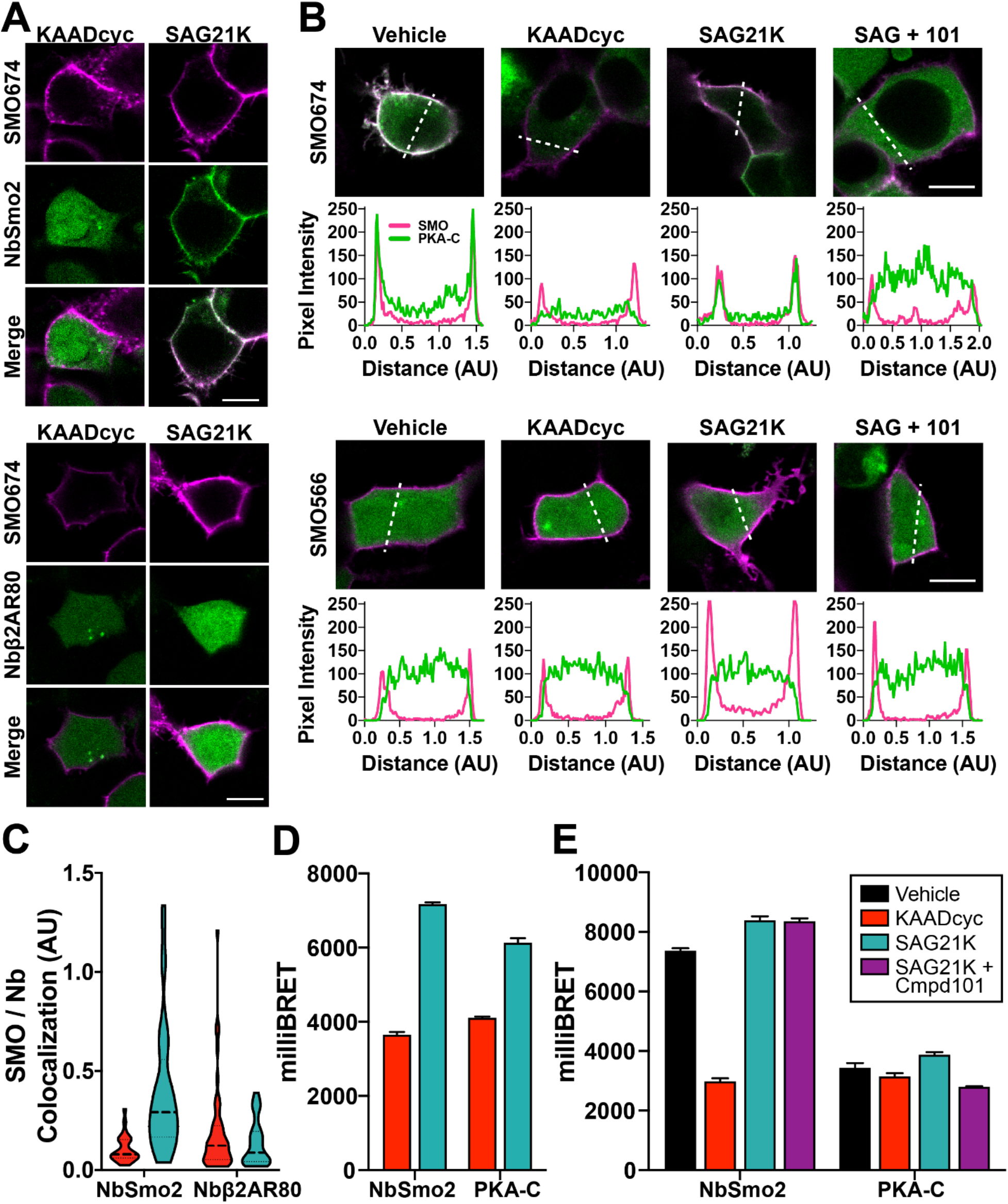
Controls for assays to look at SMO activity- and GRK2/3-dependent interactions. (**A**) HEK293 cells transfected with FLAG-tagged SMO674 (magenta) and YFP-tagged NbSmo2 or GFP-tagged Nbβ2AR80 (green) were treated with KAADcyc or SAG21k and imaged as described in **Figure 5B**. (**B**) Line scan analysis of images from cells expressing SMO674 **(Figure 5B)** or SMO566. Cells were treated as described in **Figure 5B**. (**C**) Quantification of colocalization between SMO and NbSmo2 or Nbβ2AR80 for the experiment in (A) (see “Methods”). (**D**) Raw (non-normalized) BRET ratios from **Figure 5A**. (**E**) Raw (non-normalized) BRET ratios from **Figure 5D**. See Supplemental Table 1 for statistical analysis.

**Figure 6—figure supplement 1:**
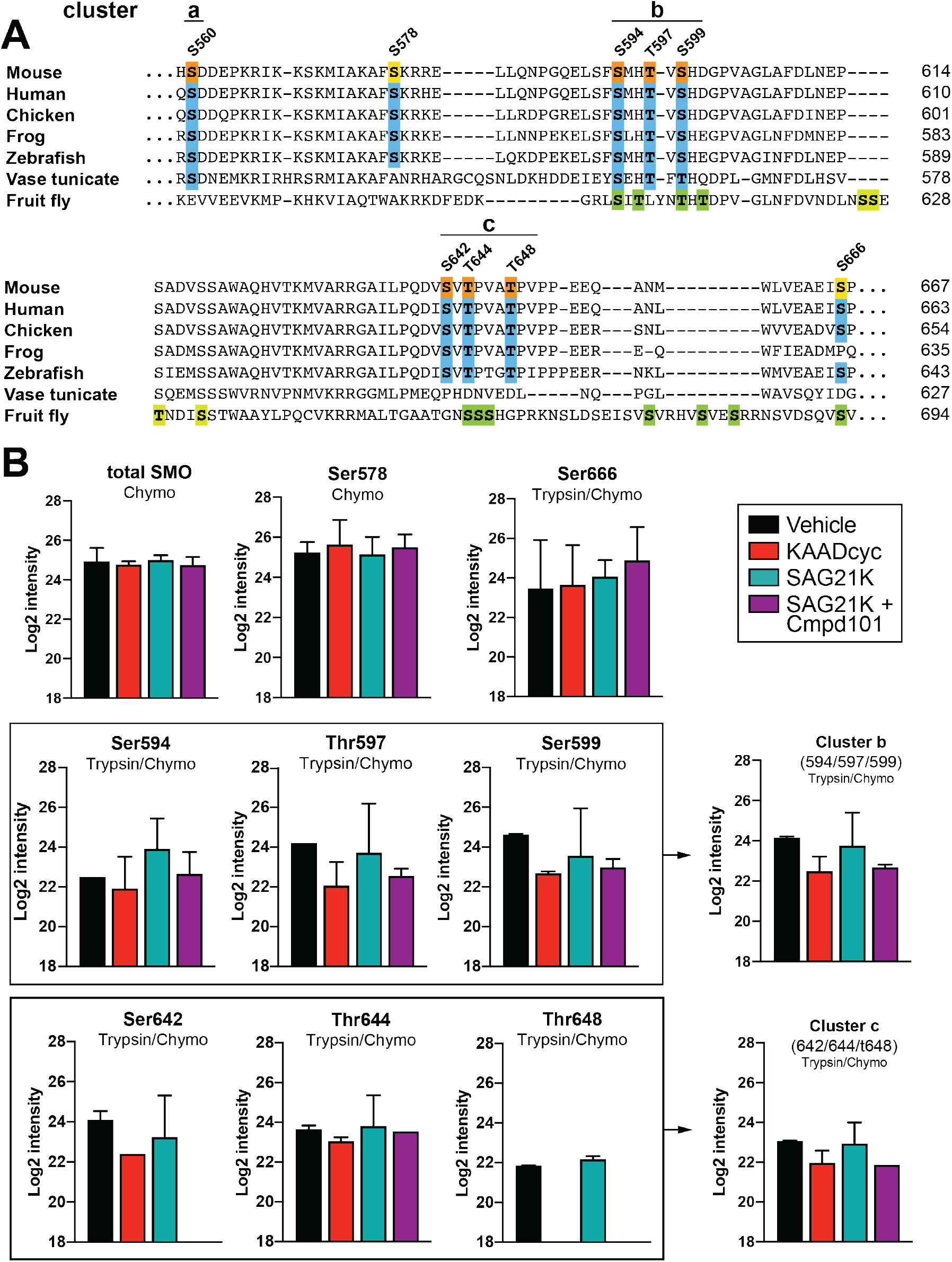
Full sequence alignments and quantification info for mass spectrometry. (**A**) Full alignment of SMO from **Figure 6B**, noting conservation of GRK2/3 among vertebrates, insects and basal metazoans. Interestingly, some GRK2/3 phosphosites in mouse SMO are charged (D or E) residues in dSmo (i.e., S560), and vice versa (i.e. D601), consistent with the importance of negative charges at those positions. dSmo phosphorylation sites are from (Maier et al., 2014), in which sites verified as GRK-dependent are indicated in dark green, while sites that might be GRK-dependent but were not covered during their mass spectrometry are indicated in light green. (**B**) Quantification of additional SMO and GRK2/3 activity-dependent phosphorylation sites and phosphorylation clusters from **Figure 6B,C**, as well as quantification of SMO following digests with the indicated proteases (“chymo” = chymotrypsin) used in our analysis (see Methods). Phosphorylation was not detected at S642 with SAG21k/Cmpd101 treatment, or at T648 with KAADcyc or SAG21k/Cmpd101 treatment, suggesting that any phosphorylation under these conditions lies below the limit of detection. Unambiguous phosphosite localization on peptides with multiple candidate phosphorylation sites in close proximity can be challenging by mass spectrometry (Potel et al., 2018) (see Methods). For these reasons, our quantification, and subsequent mutational analysis, focuses on clusters of residues rather than individual sites. n = 3 biological and 3 technical replicates per condition.

**Supplemental Table 1:** Tests of statistical significance for all figures

| Figure. | Condition 1 | Condition 2 | Significant? | p-value (Student's t-test) |
| --- | --- | --- | --- | --- |
| 1B | PKAC: Vehicle | PKAC + SMO: Vehicle | yes | 0.005967 |
| 1B | PKA-C + SMO: Vehicle | PKA-C + SMO: KAADcyc | yes | 0.010188 |
| 1B | PKA-C + M2AChR: Vehicle | PKA-C + M2AChR: Carbachol | no | 0.353043 |
| 1-suppl1C | SMO657: ShhN | SMOdelta561-657: ShhN | yes | .000007 |
| 1-suppl1D | wild type cells: M2AChR + fsk: vehicle | wild-type cells: M2AChR + fsk: Carbachol | yes | 0.002942 |
| 1C | PKA-C: Vehicle | PKA-C + SMO: Vehicle | yes | 0.000443 |
| 1C | PKA-C + SMO: Vehicle | PKA-C + SMO: KAADcyc | yes | 0.035508 |
| 1-suppl1D | Gα-null cells: M2AChR: fsk + vehicle | Gα-null cells: M2AChR: fsk + carbachol | no | 0.75318 |
| 3A | SMO + PKA-C | SMO + βarrestin1 | yes | <0.000001 |
| 3B | 657: PKA-C | delta561-657: PKA-C | yes | 0.00007 |
| 3B | 566: βarrestin1 | 566: PKA-C | no | 0.072522 |
| 3C | full-length: βarrestin1 | full-length: PKA-C | yes | <0.000001 |
| 3C | 657: βarrestin1 | 657: PKA-C | yes | 0.000005 |
| 3C | 614: βarrestin1 | 614: PKA-C | yes | 0.000002 |
| 3C | 574: βarrestin1 | 574: PKA-C | yes | 0.002667 |
| 3C | 566: βarrestin1 | 566: PKA-C | no | 0.445983 |
| 3-suppl1B | SMO566 | PTCH1 | no | 0.915218 |
| 3-suppl1E | SMO657: βarrestin1-YFP | SMO657: NbSmo2-YFP | yes | <0.000001 |
| 3-suppl1E | SMO657: βarrestin1-YFP | SMO657: PKA-C-YFP | yes | 0.000004 |
| 3D | SmoCT: βarrestin1 | SmoCT: PKA-C | yes | 0.000015 |
| 4B | PKA-C | PKA-R | yes | 0.00012 |
| 4C | PKA-C-YFP: vehicle | PKA-C-(no tag) + PKA-R-YFP: vehicle | yes | 0.000085 |
| 4C | PKA-C-YFP + PKA-R (no tag): vehicle | PKA-C-YFP + PKA-R (no tag): fsk | yes | 0.010792 |
| 4E | WT PKA-C | H87Q / W196R | no | 0.682648 |
| 4E | WT PKA-C | L206R | yes | 0.027339 |
| 4E | WT PKA-C | K73H | yes | 0.007473 |
| 4F | WT PKA-C | delta1-24 | yes | 0.000368 |
| 4F | WT PKA-C | delta1-39 | yes | 0.000095 |
| 5A | NbSmo2: SAG21k | NbSmo2: KAADcyc | yes | 0.000002 |
| 5A | PKA-C: SAG21k | PKA-C: KAADcyc | yes | 0.000072 |
| 5C | PKA-C: SAG21k | PKA-C: KAADcyc | yes | <0.0001 |
| 5C | PKA-C: SAG21k | PKA-C: SAG21k + Cmpd101 | yes | 0.0372 |
| 5C | PKA-C: Vehicle | PKA-C: KAADcyc | yes | <0.0001 |
| 5-suppl1C | NbSmo2: KAADcyc | NbSmo2: SAG21k | yes | <0.0001 |
| 5-suppl1C | Nbβ2AR80: KAADcyc | Nbβ2AR80: SAG21k | no | 0.1977 |
| 5-suppl1C | NbSmo2: KAADcyc | Nbβ2AR80: KAADcyc | no | 0.0747 |
| 5D | NbSmo2: SAG21k | NbSmo2: SAG21k + Cmpd101 | no | 0.896283 |
| 5D | PKA-C: SAG21k | PKA-C: SAG21k + Cmpd101 | yes | 0.000213 |
| 5E | PKA-C + SMO674: Vehicle | PKA-C + SMO674: Cmpd101 | yes | 0.012301 |
| 6C | S560: Vehicle | S560: KAADcyc | yes | <0.001 |
| 6C | S560: SAG | S560: SAG21K + Cmpd101 | yes | <0.01 |
| 6D | SMO657: PKA-C | SMO657Ala: PKA-C | yes | 0.00249 |
| 7A | SMO: ShhN | SMOdelta570-581; ShhN | yes | 0.000252 |
| 7B | SMO657: PKA-C | SMO657(delta570-581): PKA-C | yes | <0.000001 |
| 7D | SMO-Nbβ2AR80: Vehicle | SMO-Nbβ2AR80: ShhN | yes | 0.000216 |
| 7D | SMO-NbSmo2: Vehicle | SMO-NbSmo2: ShhN | no | 0.453546 |
| 7F | SMO657: ShhN | SMO657-Ala: ShhN | yes | 0.000198 |

